# DASH: A versatile and high-capacity gene stacking system for plant synthetic biology

**DOI:** 10.1101/2025.03.04.641457

**Authors:** Chengsong Zhao, Anna N. Stepanova, Jose M. Alonso

**Affiliations:** Department of Plant and Microbial Biology, Genetics and Genomics Academy, North Carolina State University, Raleigh, North Carolina, United States of America

**Keywords:** DNA assembly, Gene Stacking, Recombineering, Golden Braid, *PhiC31* Integrase

## Abstract

DNA assembly systems based on the Golden Gate method are popular in synthetic biology but have several limitations: small insert size, incompatibility with other cloning platforms, DNA domestication requirement, generation of fusion scars, and lack of post-assembly modification. To address these obstacles, we present the DASH assembly toolset, which combines features of Golden Gate-based cloning, recombineering, and site-specific recombinase systems. We developed (1) a set of donor vectors based on the GoldenBraid platform, (2) an acceptor vector derived from the plant transformation-competent artificial chromosome (TAC) vector, pYLTAC17, and (3) a re-engineered recombineering-ready *E. coli* strain, CZ105, based on SW105. The initial assembly steps are carried out using the donor vectors following standard GoldenBraid assembly procedures. Importantly, existing parts and transcriptional units created using compatible Golden Gate-based systems can be transferred to the DASH donor vectors using standard single-tube restriction/ligation reactions. The cargo DNA from a DASH donor vector is then efficiently transferred *in vivo* in *E. coli* into the acceptor vector by the sequential action of a rhamnose-inducible phage-derived PhiC31 integrase and arabinose-inducible yeast-derived Flippase (FLP) recombinase using CZ105. Furthermore, recombineering-based post-assembly modification, including the removal of undesirable scars, is greatly simplified. To demonstrate the utility of the DASH system, a 116 kb DNA construct harboring a 97 kb cargo consisting of 35 transcriptional units was generated. One of the CDSs in the final assembly was replaced through recombineering, and the *in planta* functionality of the entire construct was tested in both transient and stable transformants.

## Introduction

The development of DNA assembly methods, including Gibson assembly (Gibson *et al*., 2009), Gateway cloning and In-Fusion (Hartley, 2000; Park *et al*., 2015), among others (Kirchmaier *et al*., 2013; Sasaki *et al*., 2004), has greatly facilitated the widespread adoption of synthetic biology approaches. Among all DNA assembly methods, Golden Gate (Engler *et al*., 2008, 2009) and its variants, such as GoldenBraid (Dusek *et al*., 2020; Sarrion-Perdigones *et al*., 2011, 2013; Vazquez-Vilar *et al*., 2017), MoClo (Weber *et al*., 2011), Mobius (Andreou and Nakayama, 2018), Loop (Pollak *et al*., 2019, 2020), and others (Blázquez *et al*., 2023; Chamness *et al*., 2023; De Paoli *et al*., 2016; Lampropoulos *et al*., 2013; Lin and O’Callaghan, 2018; Piepers *et al*., 2023; Taylor *et al*., 2019), have gained widespread popularity due to their efficiency in combining multiple DNA fragments in a single-tube enzymatic restriction/ligation reaction. By taking advantage of Type IIS restriction enzymes, which cut DNA outside of their recognition sequence (Szybalski *et al*., 1991), Golden Gate-based cloning approaches enable the assembly of multiple DNA fragments of almost any sequence composition using a very limited number of restriction enzymes, typically from one to three. This makes these technologies particularly useful for precise and flexible manipulation of standardized DNA parts (Marillonnet and Grützner, 2020).

Despite the advantages of Golden Gate-based systems mentioned above, they also present certain challenges. For instance, a domestication step is often necessary to eliminate internal recognition sites of the enzymes used for DNA assembly, creating potentially undesired scars. The problems and inconveniences associated with the domestication process are accentuated when two or more Type IIS enzymes are employed to allow for additional DNA assembly cycles required when combining multiple transcriptional units (Bird *et al*., 2022; Marillonnet and Grützner, 2020). By using DNA methylases that target sequences partially overlapping with the recognition sites of a Type IIS enzyme, the MetClo system can utilize a single Type IIS restriction enzyme even during multi-step assemblies (Lin and O’Callaghan, 2018). This is achieved by engineering the flanking sequences of some of the Type IIS recognition sites to overlap with the target sequence of a specific DNA methylase (Lin and O’Callaghan, 2018). These sites’ methylation status can then be switched on or off by propagating the MetClo plasmids in *E. coli* strains expressing or not the corresponding methylase. Although the strategy adopted by MetClo bypasses to some degree the need for domesticating DNA fragments for multiple Type IIS enzymes, it has the inconvenience of switching between different *E. coli* strains.

In addition to the domestication process, adopting standardized syntax for the exchange of DNA parts also represents a source of scars, as each DNA part type (promoter, coding sequence, terminator, etc.) is flanked by an agreed-upon four-nucleotide fixed sequence aka a “part’s code”[26]. In this case, the scars created by following the grammar rules are located at each of the DNA part’s junctions. To reduce the potential impact of these junction scars, the Start-Stop Assembly technology uses the Type II enzyme with three-instead of a four-nucleotide overhang at each part’s flanks (Taylor *et al*., 2019). Even though this approach can eliminate the scars at the start and stop codons, scars are still formed at the junction of the other standardized DNA parts. Thus, none of the Golden Gate-based cloning systems can be considered entirely scar-free.

In addition to the use of common three-or four-nucleotide codes, the compatibility between different Golden Gate-based systems also depends on the Type IIS enzymes used (Bird *et al*., 2022). Thus, for example, even though the Type IIS restriction enzyme, *BsaI*, is often used to generate transcriptional units (Blázquez *et al*., 2023), its use is not universal among Golden Gate-based systems (De Paoli *et al*., 2016). Moreover, when selecting a second or third Type IIS restriction enzyme to enable additional cycles of DNA assembly, the choice of enzymes across Golden Gate cloning systems becomes even more inconsistent. For example, GoldenBraid 3.0 uses *BsaI* and *BsmBI,* while MoClo leverages *BsaI* and *BpiI,* and Loop relies on *BsaI* and *SapI* (Pollak *et al*., 2019; Vazquez-Vilar *et al*., 2017; Weber *et al*., 2011).

Another common challenge of most Golden Gate-based systems is the lack of efficient post-assembly modification capabilities. This means that any modification, such as replacing a promoter sequence to alter the expression levels of a gene in a multigene construct, necessitates creating a new transcriptional unit with the alternative promoter and reassembling the entire genetic circuit, often from scratch. This represents an important barrier for the “design-build-test-learn” (DBTL) cycle characteristic of synthetic biology approaches, justifying the interest in developing efficient and flexible post-assembly-modifiable, Type IIS-compatible cloning and assembly systems. Homology-based recombination approaches such as recombineering (Alonso and Stepanova, 2015; Brumos *et al*., 2020; Lee *et al*., 2001; Yu *et al*., 2000; Zhang *et al*., 1998) are especially suited for this type of post-assembly modifications as they allow for the insertion, deletion, and replacement of basically any sequence in a plasmid by using the recombineering molecular machinery engineered into *E. coli* strains such as SW105 (Warming, 2005). In this recombineering strain, three lambda-Red phage genes, *exo*, *beta,* and *gam,* are integrated into the bacterial genome and expressed under the strict control of the temperature-inducible λ*p*L/*p*R-cI857 system. The induction of these three proteins leads to very high levels of recombination between the linear donor DNA and its target, regardless of whether the target gene is located in a plasmid or within the bacterial genome. Importantly, this recombination is directed by as little as 40 nt of homology arms, i.e., sequences with 100% identity between the flanks of the donor molecule and the target DNA (Yu *et al*., 2000).

The final limitation of Golden Gate-based systems is the substantial decrease in assembly efficiency as the size of DNA fragments increases, which is further compounded by the typically limited cargo capacity of most plasmids used in these platforms. In fact, the cargo capacity of the plasmid typically used in Golden Gate-based systems rarely exceeds 25-50 kb (Pollak *et al*., 2019). The MetClo system discussed above utilizes a high-capacity vector with a p15a or F replication origin that confers a low plasmid copy number and an increased cargo capacity (Lin and O’Callaghan, 2018). However, simply increasing the cargo capacity of a plasmid does not address the limitations of the Golden Gate-based systems to operate efficiently with large DNA molecules, with the high-capacity MetClo system still suffering from reduced cloning efficiency when larger fragments are being subcloned (Lin and O’Callaghan, 2018). Therefore, to effectively handle larger DNA fragments, Golden Gate-based systems need not only to increase the cargo capacity of the plasmids used, but also to improve efficiency in assembling these larger DNA fragments. One approach to addressing the low assembly efficiency of large DNA fragments is using site-specific recombinases such as Cre, FLP, PhiC31, or Bxb1 (Andrews *et al*., 1985; Kim *et al*., 2003; Sternberg *et al*., 1986; Thorpe and Smith, 1998). These enzymes efficiently catalyze the integration and excision of large circular DNA molecules, overcoming the limitations of other *in vivo* recombination-based systems, such as recombineering and jStack, which are constrained by the low efficiency of introducing large linear DNA molecules into cells (Shih *et al*., 2016).

One elegant example of the *in vivo* use of site-specific recombinases for the generation of plant transformation-ready constructs is the GAANTRY (<u>g</u>ene <u>a</u>ssembly in *<u>A</u>grobacterium* by <u>n</u>ucleic acid transfer using recombinase technolog<u>y</u>) gene stacking system (Collier *et al*., 2018). This system consists of an *Agrobacterium* strain (ArPORT1) carrying a disarmed virulent pRi plasmid engineered with a recombinase P site, two donor plasmids (B donor and P donor) and two helper vectors (B helper and P helper) with recognition sites and the corresponding recombinases (A118 and ParA or TP901-1 and ParA), respectively. The transient expression of the A118 recombinase results in the integration of the whole B donor vector into the P site of the pRi plasmid, whereas the ParA recombinase catalyzes the excision of the backbone sequences of the inserted B donor vector. The DNA cargo integration process can be repeated many times by alternating the transformation of the B donor plus helper in one cycle and the P donor plus helper vectors in the next to generate large multigene DNA constructs. Importantly, the efficiency of the integration process is close to 100%, and the resulting *Agrobacterium* strain can be directly used to transform plants (Collier *et al*., 2018). One limitation associated with the use of the *Agrobacterium* non-binary pRi acceptor plasmid is that any post-assembly modifications remain challenging (Bian *et al*., 2022; Collier *et al*., 2018). Due to the sequential nature of gene stacking, introducing a modification in the final DNA construct requires not only generating a new DNA part but repeating all of the corresponding gene stacking steps.

Another exciting example of a highly efficient, *in vivo* recombinase-based stacking system is the TGSII system (TransGene Stacking II) that utilizes the recombinase Cre and a combination of wild-type and self-compatible *loxP* mutant sites to carry out the plasmid integration and backbone excision reactions (Zhu *et al*., 2017). In contrast with the GAANTRY system, however, the integration of a second cargo DNA fragment requires additional steps, including the extraction, purification, and digestion of the resulting plasmid mixture with the *I-SceI* or *PI-SceI* homing endonuclease in order to eliminate undesired plasmid species and isolate the intended acceptor vector-cargo DNA complex (Zhu *et al*., 2017). The addition of these *in vitro* steps makes this method somewhat more cumbersome than the GAANTRY system, where the acceptor vector remains in the *Agrobacterium* cells throughout the stacking process. Importantly, like the GAANTRY system, the TGSII is not fully integrated with the Golden Gate-based system and, by itself, lacks post-assembly modification capabilities.

To overcome some of the technical barriers often found in Golden Gate-based systems, we present an innovative assembly and stacking system named DASH (<u>D</u>NA <u>a</u>ssembly and <u>s</u>tacking <u>h</u>ybrid). This system takes advantage of the common restriction enzyme, *BsaI,* to enable compatibility with standardized DNA parts and transcriptional units from other Golden Gate-based cloning systems. The DASH system combines the reiterative assembly capabilities of the GoldenBraid system, the high cargo capacity of the transformation-competent bacterial artificial chromosome (TAC) single-copy binary vector pYLTAC17, the high *in vivo* integration/excision efficiency of the PhiC31 and FLP recombinases, and the precision of the genome editing recombineering technology at introducing any sequence changes in DNA molecules of any size. Importantly, the gene stacking capabilities of the DASH platform can be used to overcome the size limitations not only of DNA constructs assembled using the DASH and GoldenBraid systems but also those of transcriptional units and modules generated in other Golden Gate-based cloning systems that use the *BsaI* enzyme in their assembly, such as Loop, MoClo, Mobius, and Start-Stop (Figure 1)(Andreou and Nakayama, 2018; Pollak *et al*., 2019; Taylor *et al*., 2019; Weber *et al*., 2011). Finally, the TAC acceptor vector in DASH has the capacity to stably maintain a large DNA fragment of up to 300 kb (Tao, 1998), while the recombineering machinery is available for post-assembly modification, including insertion, deletion, and replacement of any sequence precisely and efficiently.

**Figure 1.**
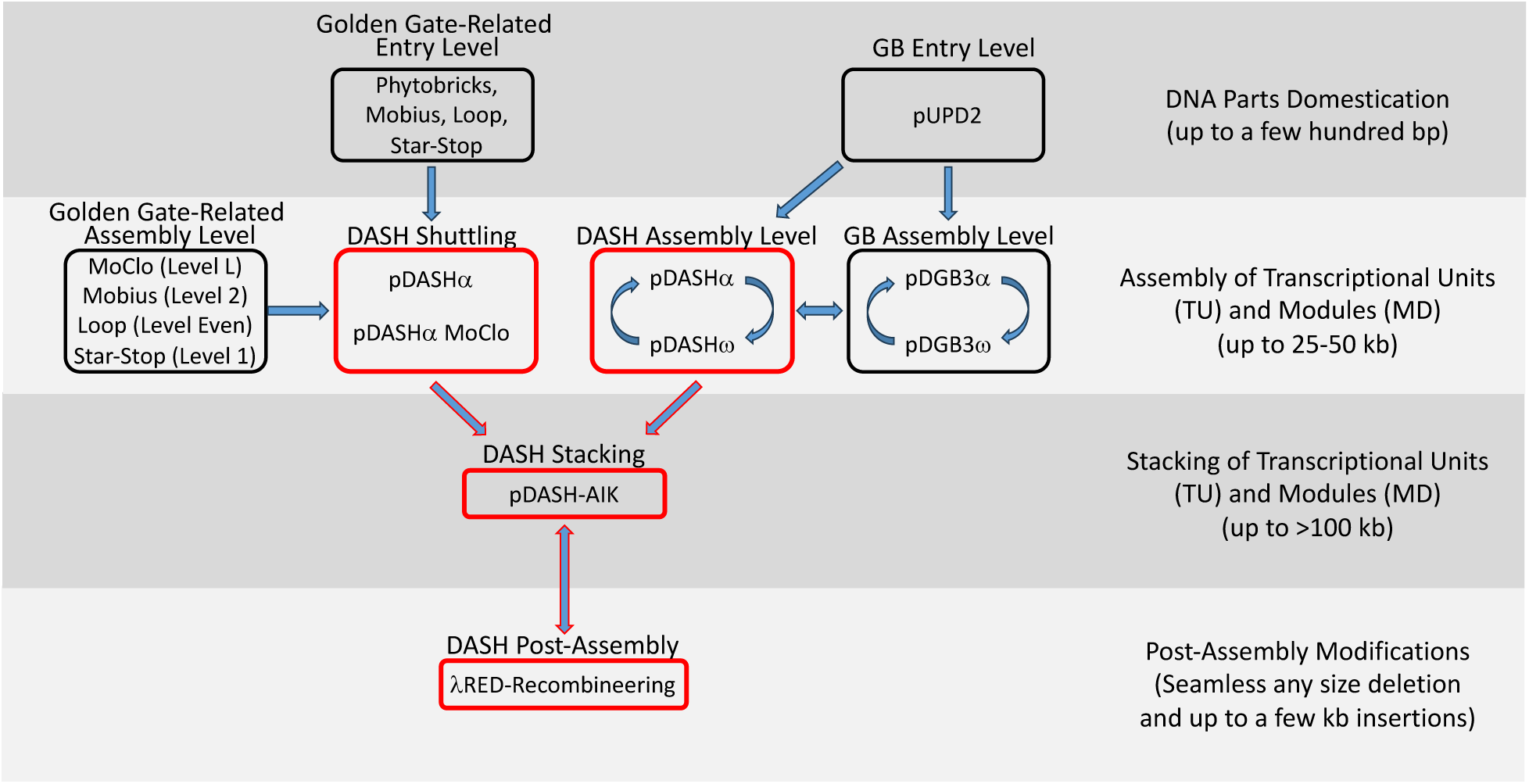
Compatibility of the DASH assembly and gene stacking system with other common DNA assembly platforms. Like all other systems based on Golden Gate, DASH can be utilized to assemble domesticated DNA parts into transcriptional units (TUs). Once these TUs are in a DASH vector, they can be further combined into modules (MDs) in a typical reiterative binary assembly procedure characteristic of the GoldenBraid (GB) system as long as all DNA parts have been domesticated for both *BsaI* and *BsmBI*. Thus, the DNA assembly components of the DASH system are fully compatible with GoldenBraid parts, TUs, and MDs. Parts domesticated for other systems that also use *BsaI* but not *BsmBI*, such as Mobius, Loop, and MoClo, can be assembled into TUs using the pDASH alpha vectors to facilitate their subsequent stacking in the pDASH-AIK destination vector. Likewise, DNA constructs in Level Even of Loop, Level 2 of Mobius, and Level 1 of Star-Stop can also be directly transferred to a pDASH alpha vector using the *BsaI* enzyme. The system’s compatibility can be further enhanced by generating shuttle vectors such as the pDASHa MoClo, which allows the transfer of DNA constructs from MoClo Level M vector into the DASH system. Once a DNA construct is in one of the pDASH alpha vectors, it can be directly used in a stacking experiment and transferred to the pDASH-AIK destination vector or, if lacking the *BsmBI* recognition sites, combined with other DNA constructs already in the DASH system. Thus, the DASH system enables the stacking of DNA constructs created using various assembly method platforms and overcomes the size limit for GoldenBraid, Loop, Mobius, MoClo, and Start-Stop systems. Moreover, recombineering can be utilized to create any post-assembly modifications after the DNA constructs are stacked in the pDASH-AIK destination vector.

Notably, the DASH system has been designed for high efficiency and experimental simplicity. Thus, all of the initial assembly steps to generate transcriptional units and small multigenic constructs take advantage of the single tube restriction/ligation reaction characteristic of Golden Gate-based systems. Single genes or multigenic constructs generated using the DASH system or a variety of Golden Gate-based platforms can easily and efficiently be stacked into a high-capacity acceptor vector (Figure 1). This is achieved by simply electroporating the cargo-carrying DASH donor vector into the SW105-derived *E. coli* strain, CZ105, containing the pDASH-AIK acceptor vector and activating different recombinases (PhiC31 and FLP) *in vivo* by growing the cells in the presence of different sugar inducers (rhamnose or arabinose, respectively). The selection of the cargo-containing acceptor vector is also done *in vivo* using antibiotics, while the removal of the donor plasmid and the corresponding cargoless backbone can be facilitated by the *SacB* contra-selectable marker. Finally, the temperature-inducible recombineering system integrated into the CZ105 cells further simplifies the post-assembly modification process, minimizing the need for the *in vitro* manipulation of large DNA molecules (Figure 1). We demonstrated the practicality of the DASH system by generating a 35-gene cassette with a total DNA insert of over 97 kb and we validated the functionality of this large construct in *Nicotiana benthamiana* transient expression assays and stable *Arabidopsis thaliana* transformants. Finally, we also confirmed the post-assembly capabilities of the system by replacing a transcriptional unit containing *superfolder GFP* (*sfGFP*) with the *RPSL-Amp* cassette in the final ∼97 kb construct cargo by leveraging the recombineering feature of the DASH system.

## Results

### Design of the DNA Assembly and Stacking Hybrid (DASH) system

The core DASH system consists of three basic components: the SW105-derived *E. coli* strain (CZ105a), a set of four donor vectors derived from the GoldenBraid assembly system (pDASH-DII-α1, pDASH-DI-α2, pDASH-DI-ω1, and pDASH-DII-ω2), and the pYLTAC17-derived acceptor vector (pDASH-AIK) (Figure 2A). In addition to these basic components, we have developed several additional vectors to expand the capabilities of the system. Thus, for example, pDASH-DII-α1 (MoClo) and pDASH-DI-α2 (MoClo) were developed to illustrate how simple modification of the core donor plasmids could be used to increase the compatibility between DASH and other Type IIS-based DNA assembly systems. Similarly, to facilitate the removal of the donor plasmids and their derived backbone after the integration of the cargo into the acceptor vector, a second set of donor vectors containing the *SacB* counter-selectable marker was also created (Figure 2B). In addition, a second acceptor vector, pDASH-AIIK, was generated to enable the initiation of the stacking process with a DII-type donor vectors instead of a DI.

**Figure 2.**
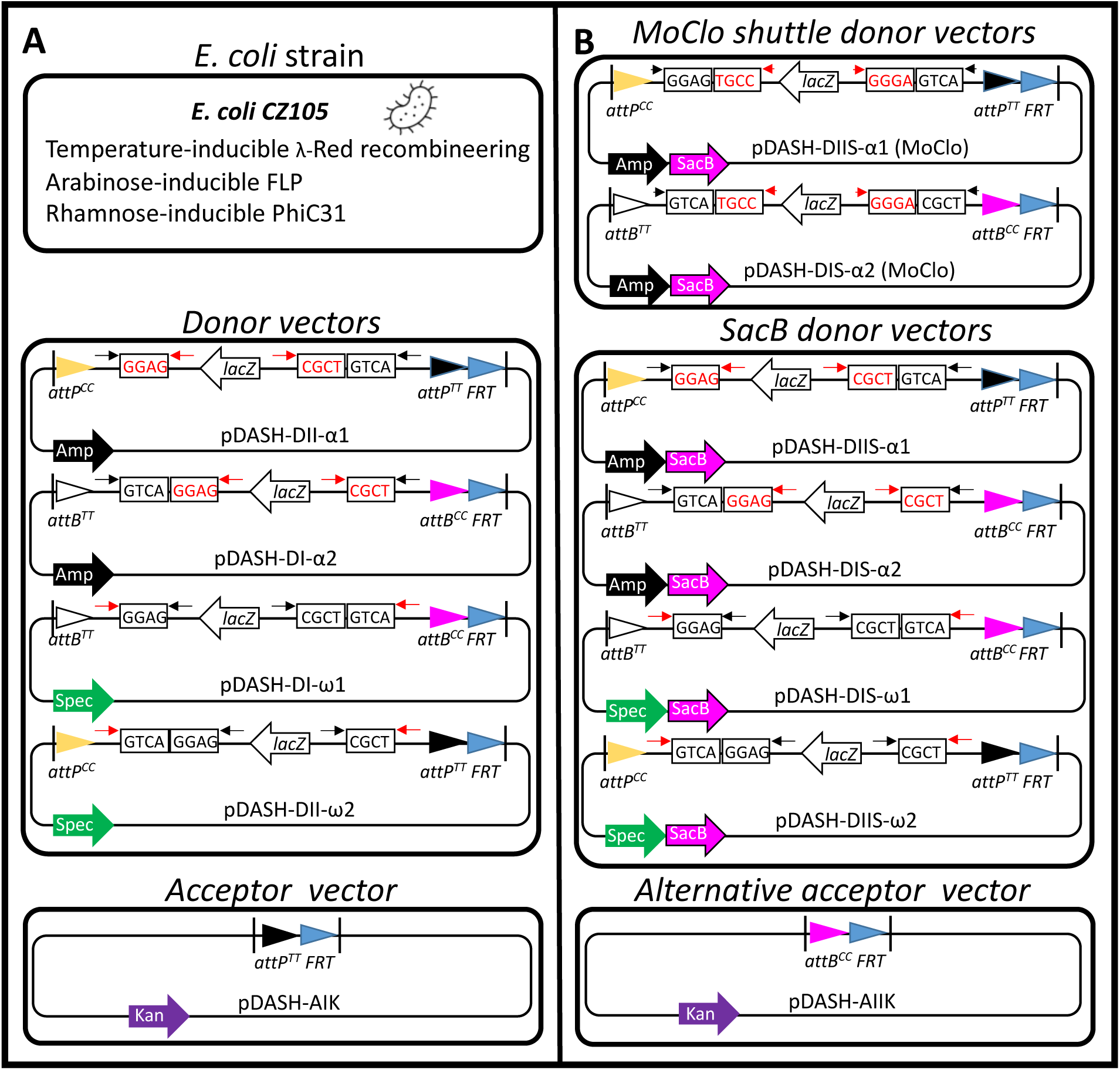
Key components of the DASH system. (A) DASH system is composed of three essential elements. 1) The CZ105a strain of *E. coli* carries in its genome the temperature-inducible lambda-Red recombineering system, an arabinose-inducible *FLP* recombinase, and a rhamnose-inducible *PhiC31* integrase (A, top panel). 2) Four donor vectors derived from the pDGB GoldenBraid plasmids that incorporate the recognition sites for the PhiC31 integrase and the FLP recombinase. Using two sets of orthogonal *attB-attP* sites allows the reiterative insertion of the donor vectors into the acceptor vector following a DI➔DII➔DI➔DII, etc. arrangement. The presence of *FRT* sites facilitates the removal of the donor vector backbone upon integration (A, middle panel). 3) A single acceptor vector derived from the TAC pYLTAC17 vector but containing an *attP^TT^* site to facilitate the integration of the first donor vector and the *FRT* site to allow for the elimination of the donor vector backbone (A, bottom panel). (B) To improve the system’s compatibility and aid in removing the donor vector-derived sequences, we created two additional sets of vectors. The MoClo shuttle donor vectors are modifications of the standard pDASH alpha vectors, designed so that after digestion with the *BsaI* enzyme, the resulting overhangs are compatible with those used in the MoClo Level M vectors (B, top panel). In addition, the *SacB* counter-selectable marker gene was added to all four pDASH donor vectors to facilitate their elimination from *E. coli* after the integration of their cargo into the acceptor vector (B, middle panel). The second acceptor vector with the *attB^CC^* site was also produced to facilitate the initiation of the stacking process with the DII donor vectors (B, bottom panel). Thin black arrows mark the recognition sites for the *BsmBI* enzyme, while the thin red arrows mark the recognition sequences for the *BsaI* enzyme. Black sequences inside of rectangles indicate the overhangs created after the *BsmBI* digestion, while the red sequences in the rectangles show the overhangs generated by the *BsaI* enzyme. The different *attP, attB* and *FRT* sites are depicted as arrow heads of different colors. The different selectable marker genes (*Kan, Amp, Spec, LacZ* and *SacB*) are shown as arrows of different colors. The vertical black lines represent the left and right borders of the T-DNA.

The CZ105a DASH strain was derived from the recombineering strain SW105, which harbors a genomic insertion of a defective lambda prophage (Warming, 2005). This prophage includes the three lambda-Red recombineering-required genes: *gam, beta,* and *exo*. These genes are controlled by the temperature-sensitive cI857 repressor, which prevents or allows their expression at low (30°C) and high (42°C) temperatures, respectively (Yu *et al*., 2000). In addition, this strain also carries the arabinose-regulated araC transcription factor to drive the expression of the arabinose-inducible *FLP* recombinase gene, *PBADflpe* (Warming, 2005). We utilized the recombineering capabilities of the parental SW105 to insert the *PhiC31* sequence with a ribosomal binding site (*RBS*), *RBS-PhiC31,* downstream of the rhamnose-inducible genes *rhaA* and *rhaT* in the SW105 genome, creating the CZ105a and CZ105b DASH strains (Figure S1), respectively. This was achieved in a two-step recombineering process. First, a positive-negative selection cassette *RPSL-Amp* (Brumos *et al*., 2020) was inserted just downstream of the *rhaA* stop codon and then replaced by the *RBS-PhiC31* sequence to generate the CZ105a DASH strain in the second recombineering reaction. Using the same strategy, we also inserted the *RBS-PhiC31* sequence immediately downstream of the stop codon of *rhaT* in the SW105 strain to generate the CZ105b DASH strain.

The PhiC31 catalyzes recombination between heterotypic sites, *attP* and *attB*. This reaction is irreversible as PhiC31 alone cannot catalyze the recombination between the resulting *attR* and *attL* sequences. If the *attP* and *attB* sites are in different plasmids, their recombination will combine both molecules to generate a single hybrid plasmid. The *attP* and *attB* sequences are both palindromic with a central nonpalindromic sequence. By changing this central nonpalindromic sequence, it is possible to generate orthogonal *attP-attB* site pairs (Merrick *et al*., 2018). Thus, for example, the *attP^TT^* and the *attB^TT^*, where the ‘TT’ superscripts indicate the nonpalindromic central dinucleotide sequences, can recombine, whereas the *attP^TT^*and *attB^CC^* cannot. The orthogonality between ‘TT’ and ‘CC’ sites can be utilized for the iterative integration of donor vectors into an acceptor vector. Thus, for example, if a donor vector contains both *attB^TT^*and *attB^CC^* sites, it can be integrated into an acceptor vector containing an a*ttP^TT^* site. The resulting hybrid plasmid will contain the *attL^TT^* and *attR^TT^* inactive sites as a result of the recombination between the *attB^TT^* and *attP^TT^* sites, as well as a functional intact *attB^CC^* site. If the second donor vector has an *attP^CC^* and *attP^TT^* sites, it could be integrated into the hybrid vectors from the first integration cycle, producing inactive *attL^CC^* and *attR^CC^* and also introducing an active intact *attP^TT^* site that would be available for the next round of integration, and so on. This strategy could, in principle, be used to insert as many donor vectors as necessary into the acceptor vector, but it would result in the accumulation of not only the cargo sequences present in the donor vectors but also the vector backbones. The removal of the backbone sequences of the donor vector from the hybrid plasmids can be achieved using another site-specific recombinase, FLP, which catalyzes the recombination between two homotypic *FRT* sites (Andrews *et al*., 1985; Senecoff *et al*., 1985). If the two identical *FRT* sites are in the head-to-tail orientation in the same DNA molecule, the FLP recombinase precisely excises the sequence between the two sites, leaving a single copy of an active *FRT*. By strategically introducing *FRT* sites at selected locations in both the donor and acceptor vectors—one *FRT* in each—it becomes possible to remove the backbone of the donor vector after each *attP-attB* integration cycle.

We took advantage of the properties of the PhiC31 and FLP recombinases and their corresponding recognition sites described above to modify the existing GoldenBraid and high-cargo capacity pYLTAC17 vectors to generate the set of DASH donor and acceptor plasmids. Thus, we introduced the *attP^CC^* and *attP^TT^-FRT* sequences into the GoldenBraid pDGB3α1 and pDGB3ω2 vectors to generate the pDASH-DII-α1 and the pDASH-DII-ω2 plasmids. Similarly, the *attB^TT^* and *attB^CC^-FRT* sequences were inserted into the GoldenBraid pDGB3ω1 and pDGB3α2 vectors to generate the pDASH-DI-ω1 and the pDASH-DI-α2 plasmids, respectively. To make the pDASH-DII-α1 and pDASH-DI-α2 compatible with the acceptor vectors (pDASH-AIK and pDASH-AIIK acceptors, see below), we replaced the Kan-resistance gene in pDASH alpha vectors with a *BsaI*-domesticated Amp-resistance gene (Figure 2B, Tables S1-3). To facilitate the elimination of the pDASH donor vectors from the *E. coli* cells after each round of integration and excision, we inserted a *RBS* and the coding sequence of a *BsaI-* and *BsmBI-* domesticated *SacB* contra-selectable marker gene downstream of the *Amp* or *Spec*-resistance genes in each of the original pDASH donor vectors (Figure 2B and Table S3). To make our system compatible with the widely used MoClo assembly system, we modified the pDASH-DII-α1 and pDASH-DI-α2 donor vectors such that the overhangs generated by the *BsaI* digestion become compatible with the MoClo level M transcriptional units (Figure 2B). We call these MoClo-compatible vectors pDASH-DII-α1 (MoClo) and pDASH-DI-α2 (MoClo) (Table S3). Finally, to generate the pDASH-AIK and pDASH-AIIK acceptor vectors, we introduced the *attP^TT^-FRT* and *attB^CC^-FRT* sequences, respectively, into the large cargo capacity vector pYLTAC17 (Figure 2, Table S1).

### Functionality of the rhamnose-inducible PhiC31 expressed from a genomic locus

In *E. coli*, the *rha* regulon, which contains a rhamnose transporter gene *rhaT*, the *rhaBAD* operon for rhamnose catabolism, and the *rhaRS* regulatory operon, is responsible for L-rhamnose metabolism (Moralejo *et al*., 1993). Two promoters in this regulon, *PrhaBAD* and *PrhaT*, have been shown to be tightly controlled by rhamnose and extensively used in plasmid-based inducible systems in bacteria (Fricke *et al*., 2022; Hanko *et al*., 2020). To determine if *PhiC31* driven by these rhamnose-inducible promoters in their chromosomal context was effective at catalyzing the recombination between *attP* and *attB* sites in *E. coli*, we transformed our two DASH *E. coli* strains CZ105a and CZ105b (harboring the rhamnose-inducible *PhiC31* gene) (Figure 1S) with the pDASH-DI-ω1 donor and pDASH-AIK acceptor vectors. After growing in the presence of both spectinomycin and kanamycin in liquid LB to an OD600 of ∼0.6, the cells were pelleted and resuspended in LB containing kanamycin and 0.1% W/V L-rhamnose to induce the expression of the *PhiC31* recombinase gene. After 3h at 30° C and constant shaking, L-arabinose was added to the culture to a final concentration of 0.1%, and the cultures were incubated for another 3h at 30°C and constant shaking to induce the *FLP* expression. We observed that the CZ105a cells grew normally, while CZ105b did so slowly. After plating the induced cells in LB-kanamycin plates and incubating the cells for two days at 30°C, colony PCR was performed using the diagnostic primers P27 and P28 (Figure 3, Figure S2 and Table S1). PCR and subsequent Sanger sequencing showed that in ∼50% of colonies in the CZ105a background (10 out of 20 colonies), the *LacZ* fragment from the donor vector was correctly integrated into the acceptor vector, and the corresponding backbone was removed (Figure S2). The fidelity of the two recombination events was confirmed by Sanger sequencing of the integration junctions (Figure S3). On the other hand, 100% of colonies of CZ105b still contained the intermediate plasmid, in which the donor vector was integrated into the acceptor vector (Supplemental Sequence 1), but the excision of the donor vector backbone could not be confirmed (supplementary Figure S2). Given the poor growth of CZ105b and the unexpected difficulties with excising the backbone of the donor vector, we decided to do all the subsequent experiments with the CZ105a strain.

**Figure 3.**
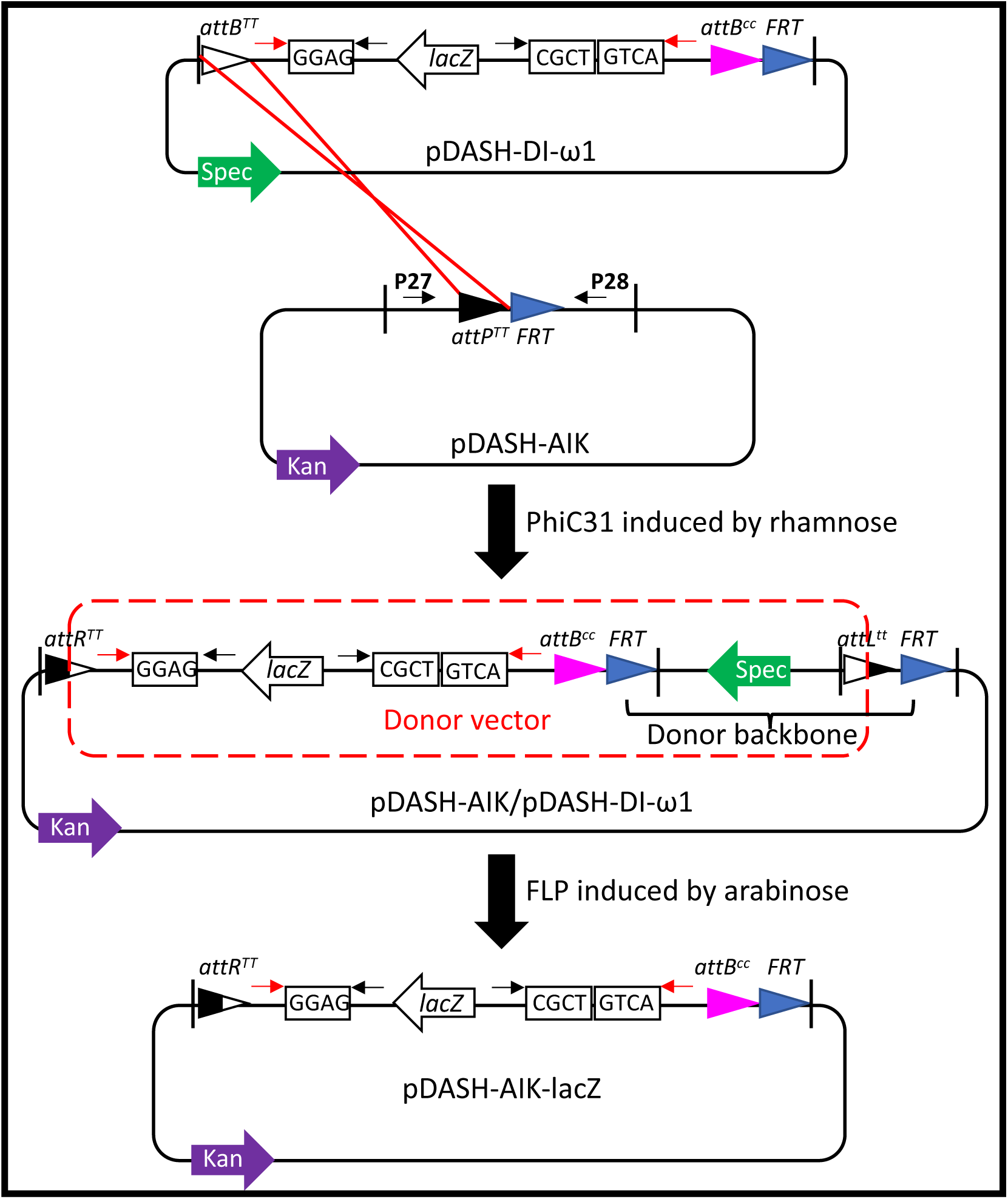
Schematic illustration of the first round of integration and excision process. Rhamnose-induced PhiC31 in the CZ105a cells catalyzes the recombination between the *attB^TT^* site in the pDASH-DI-ω1 donor vector and the *attP^TT^*site in the acceptor pDASH-AIK vector *in vivo*. The resulting intermediate hybrid plasmid pDASH-AIK/pDASH-DI-ω1 contains the whole sequences from both recombined plasmids. Arabinose-induced FLP results in the recombination between the two *FRT* sites and, therefore, the excision of the pDASH-DI-ω1 backbone sequences. A red dashed box indicates the sequences corresponding to the integrated donor vector, while the donor vector backbone is marked with a black brace. After the integration/excision process, the *attB^CC^* site transferred from the DI doner vector becomes available for a second round of integration excision using a DII vector that contains an *attP^CC^* site.

To demonstrate the system’s ability to support several rounds of integration and excision, we transformed CZ105a cells carrying the pDASH-AIK-*lacZ* from the first donor integration cycle with the pDASH-DII-ω2 donor vector. After following the same L-rhamnose and L-arabinose induction protocol described above, cells were plated in kanamycin-containing LB plates, and the integration and excision recombination products were examined by colony PCR. As with the first integration cycle, PCR screening with the diagnostic primers P27 and P28 (Figure S3 and Table S1) indicated that ∼50% of the colonies had undergone a correct second round of integration-excision recombination. As before, the fidelity of this recombination was confirmed by Sanger sequencing (Figure S3).

### Compatibility of the DASH and GoldenBraid systems

As mentioned earlier, the DASH system can be used to assemble parts and stack transcriptional units from various Golden Gate-based systems, including GoldenBraid, Loop, Mobius, MoClo, and Start-Stop. This compatibility can easily be expanded by using linkers or modified pDASH donor vectors, such as pDASH-DII-α1 (MoClo) and pDASH-DI-α2 (MoClo) (Figure 2B). To demonstrate the compatibility of the DASH system and its utility to bypass the cargo size limitation of other platforms, we started by combining eight transcriptional units in a series of binary assembly reactions using a combination of GoldenBraid and DASH vectors. These eight transcriptional units had been previously assembled in the GoldenBraid GBα1 (*35S-MCP-VPR-Tnos [TRANSCRIPTIONAL UNIT1 (TU1)] (*(Selma *et al*., 2019)*), AtU6p-sgRNA_Spy(SlDFR)_-MS2-AtU6t [TU3], 35S-dCas9_Sth_-EDLL-T35#0 [TU5], AtU6p-sgRNA_Sth(ADH1)_-MS2-AtU6t [TU7]*) or in

GBα2 (*35S-dCas9_Spy_-EDLL-Tnos [TU2]*((Selma *et al*., 2019))*, SlDFRp_Spy(SlDFR)_-3xYpet-Term35S#0 [TU4], Kan (GB0184) [TU6]* (Ahrazem *et al*., 2016)*, SlDFRp_Sth(ADH1)_-mCherry-T35S#0 [TU8]*) (Figure 4). Using standard Golden Gate assembly protocols, the *TU1* and *TU2* were combined into the GoldenBraid pDGB3 ω1 (GBω1) vector to generate the GBω1-MD1 (Module 1), the *TU3* and *TU4* were combined into the pDASH-DII-ω2 vector to form the pDASH-DII-ω2-MD2. Similarly, the *TU5* plus *TU6,* and *TU7* plus *TU8* were combined to form the pDASH-DI-ω1-MD3 and pDASH-DII-ω2-MD4, respectively. Again, using the typical braiding assembly strategy of GoldenBraid, the GBω1-MD1 and the pDASH-DII-ω2-MD2 were assembled into the pDASH-DII-α1 vector to form the pDASH-DII-α1-MD5 module, while the pDASH-DI-ω1-MD3 and pDASH-DII-ω2 –MD4 were combined into the pDASH-DI-α2 to form the pDASH-DI-α2-MD6 module. It is important to point out that all these assemblies worked with the typical high efficiency of the GoldenBraid cloning system, even though we used a mixture of vectors from both platforms, the original GoldenBraid pDGB3 and our DASH vectors. Next, we combined the pDASH-DII-α1-MD5 and pDASH-DI-α2-MD6 into the pDASH-DI-ω1 to form the pDASH-DI-ω1-MD7 (Figure 4). As we typically observe when generating constructs of over 15-20 kb in GoldenBraid pDGB3 vectors, a significant decrease in efficiency was noted in the assembly of the ∼23 kb pDASH-DI-ω1-MD7, where only ∼13% of the colonies had the expected module combination based on LacZ activity, colony PCR using the primers mCherry f and pDGB3 r, and restriction digestion diagnosis (Tables S4-5, and Figure S4A-C). We attempted to further increase the size of the pDASH-DI-ω1-MD7 by combining it with the 3.5 kb pDASH-DII-ω2-MD8 containing the *FAST* and *BASTA* (GB0023) transcriptional units to create pDASH-DI-α2-MD9. However, after screening 25 white colonies by colony PCR and the following restriction digestion, we were not able to identify any positive clones (Tables S4-5, Figure S4D-F).

**Figure 4.**
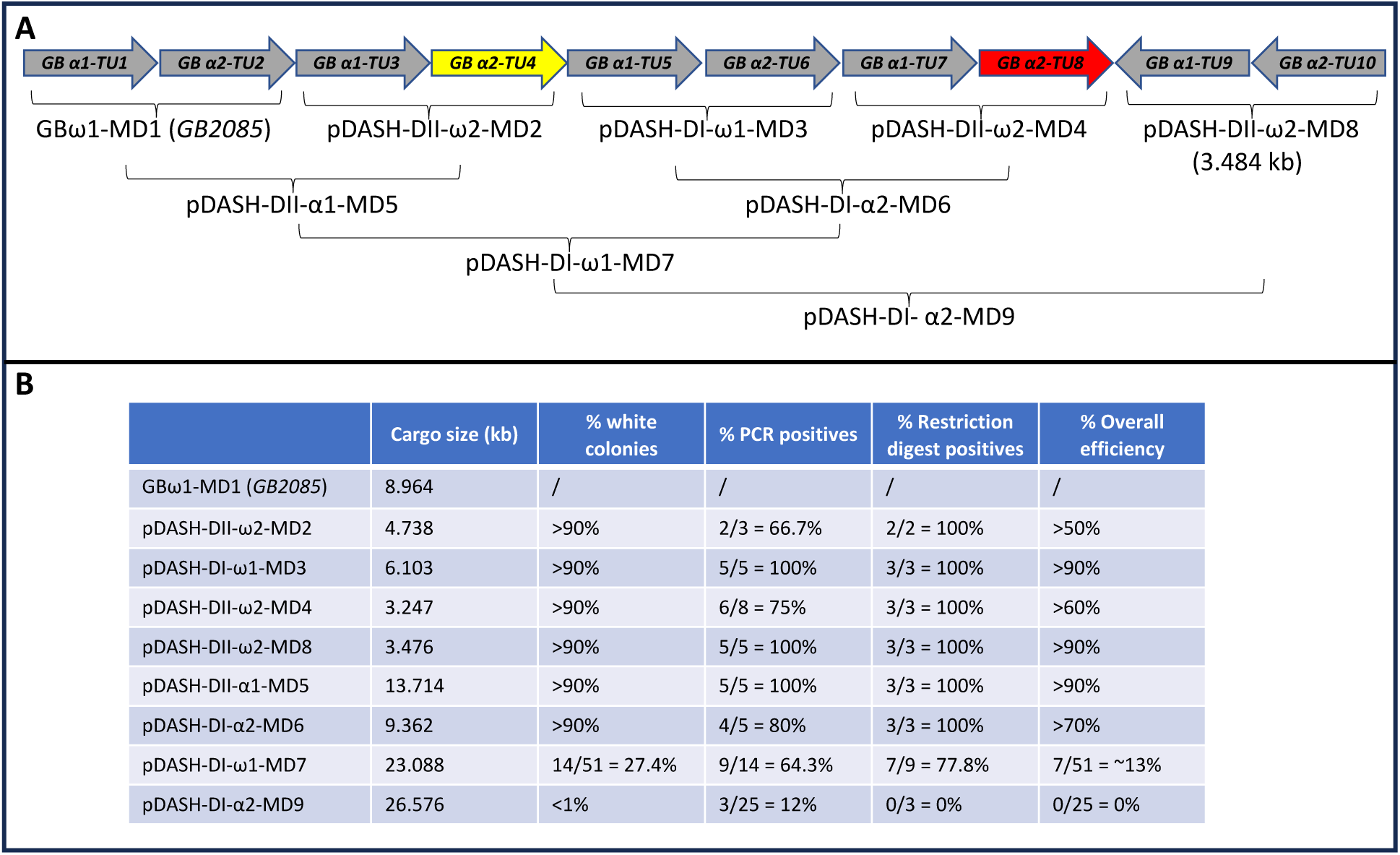
DASH donor vector assembly works seamlessly with GoldenBraid components, but it becomes inefficient for constructs larger than approximately 25 kb. (A) Transcription units (right arrows, drawing not to scale) in GoldenBraid (GB) consist of the following sequences: *35S-MCP-VPR-Tnos (GB α1-TU1), 35S-dCas9_Spy_-EDLL-Tnos(GB α2-TU2), AtU6p-sgRNA_Spy(SlDFR)_-MS2-AtU6t (GB α1-TU3), SlDFRp_Spy(SlDFR)_-3xYpet-Term35S#0 (GB α2-TU4), 35S-dCas9_Sth_-EDLL-Term35#0 (GB α1-TU5), Kan (GB0184)(GB α2-TU6), AtU6p-sgRNA_Sth(ADH1)_-MS2-AtU6t (GB α1-TU7), SlDRFp_Sth(ADH1)_-mCherry-Term35S#0 (GB α2-TU8), FAST* seed fluorescent marker *(GB α1-TU9),* and *Basta (GB0023) (GB α2-TU10).* These transcriptional units were assembled in the typical GoldenBraid pairwise fashion, as indicated by the horizontal brackets. The names of the modules (MDs) resulting from these pairwise assemblies are also shown. (B) The size of the different modules and the corresponding assembly efficiencies are shown.

### High cargo capacity of the DASH platform

The difficulties with assembling a ∼26 kb construct and the low efficiency observed in the construction of the ∼23 kb pDASH-DI-ω1-MD7 illustrate the limitations of the standard GoldenBraid assembly system in generating large, multigenic constructs. To mitigate this limitation, we have equipped the DASH system with two special features. On the one hand, the *PhiC31* recombinase present in the DASH CZ105a strain has, in principle, the capacity to combine large DNA molecules as it does during the normal life cycle of phages, where it catalyzes the integration of the ∼42 kb phage DNA into the large size bacterial host genome (Thorpe and Smith, 1998). On the other hand, the pYLTAC17-derived pDASH-AIK acceptor vector has a cargo capacity of over 100 kb (Hirose *et al*., 2015), allowing for the stable propagation of large multigenic constructs (Zhu *et al*., 2017). To test the capability of the DASH system to operate with large DNA fragments, we started by examining the efficiency of transferring the 23 kb cargo from the pDASH-DI-ω1-MD7 to the pDASH-AIK acceptor vector. Competent cells of the CZ105a strain carrying the empty pDASH-AIK were transformed with the purified pDASH-DI-ω1-MD7 plasmid. Cells were resuspended in LB containing kanamycin, and *PhiC31* and *FLP* were induced as described above. Eleven out of 20 (55%) of the randomly selected colonies showed the correct integration of the 23 kb cargo and excision of the corresponding backbone sequences of the pDASH-DI-ω1-MD7 module based on PCR using the diagnostic primers P27 2^nd^ and 35S r, mCherry f and P28 (Tables S1, 4, 5 and Figure 5). This efficiency was similar to that previously observed when integrating the much smaller *lacZ* cargo, indicating that the size of the DNA fragments transferred from the DASH donor to the DASH acceptor vectors did not affect the efficiency of the integration/excision process. To further test the potential influence of the cargo size in the DASH acceptor vector on the integration efficiency, we carried out six additional rounds of integration/excision (Figure 5), alternating the incorporation of a unique-sequence single transcriptional unit and a module of eight transcriptional units of the pDASH-DI-ω1-MD7. Thus, in the subsequent rounds of integration excision, the donor vectors pDASH-DII-α1 (containing *LacZ*), pDASH-DI-ω1-MD7, pDASH-DII-α1-*FAST*, pDASH-DI-ω1-MD7, pDASH-DIIS-α1-*sfGFP*, and pDASH-DI-ω1-MD7 were used to generate the 35-transcriptional unit pDASH-AIK-Big construct (Figure 5). Importantly, the integration/excision efficiency remained constant at around 50% (Figure 5), confirming that neither the size of the donor nor of the acceptor vectors has a significant effect on the efficiency of the stacking system. The fidelity of the integration excision process was confirmed by diagnostic PCR using five pairs of primers (Table S5) and whole-plasmid sequencing (Data S1).

**Figure 5.**
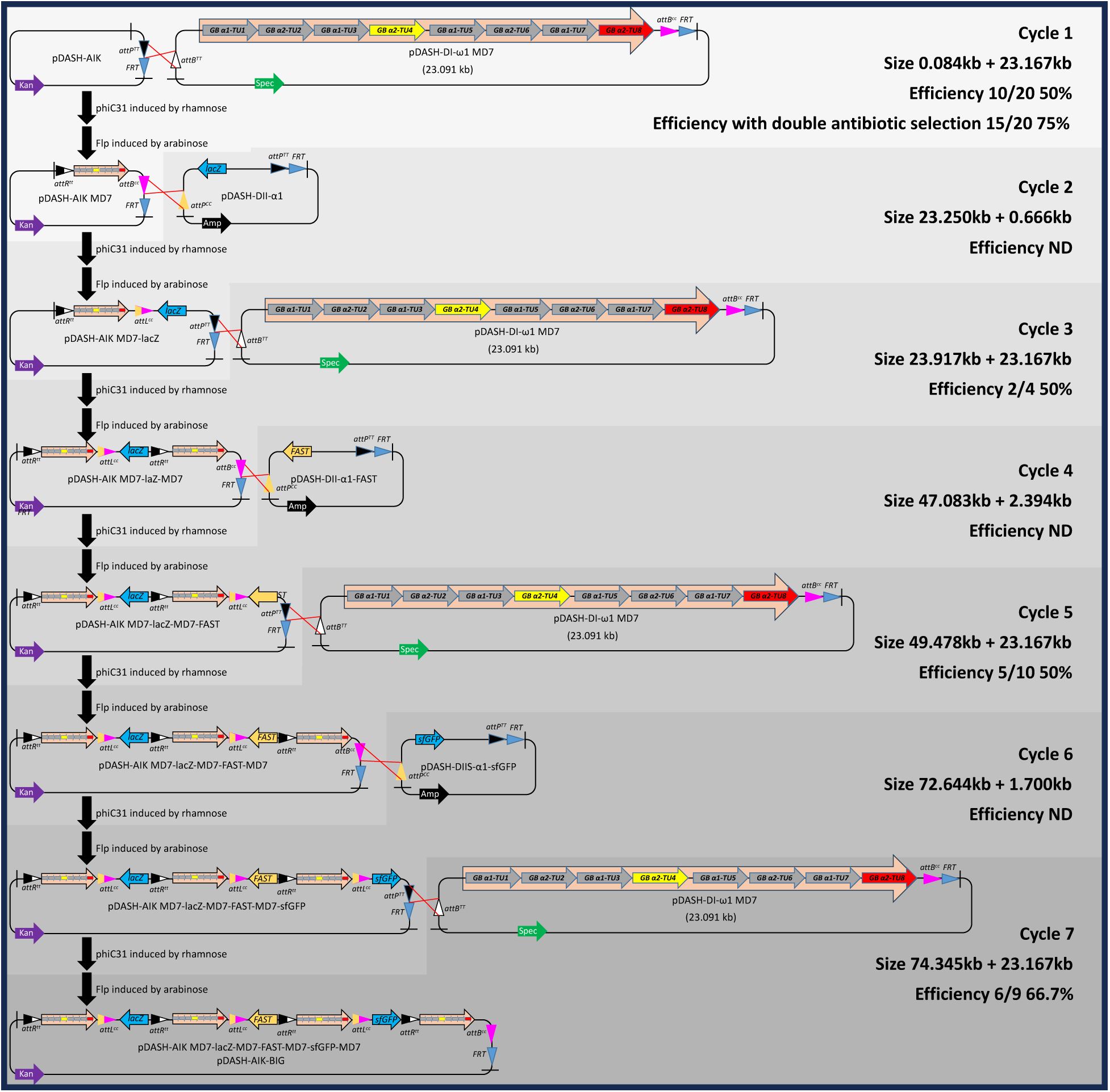
The DASH system can efficiently generate large gene stacks of over 90 kb. Schematic representation of the seven consecutive integration/excision reactions used to stack 35 transcriptional units in the final destination plasmid. The PhiC31 recombination sites involved in each stacking cycle between a donor and acceptor vector are shown, as well as the architectures of the resulting plasmids. Transcriptional units are shown as broad arrows, while large modules are shown as large orange arrows containing transcriptional unit arrows. Triangles depict recombination sites for either PhiC31 or FLP. The cargo size of each of the vectors, as well as the efficiency of each integration/excision cycle, are also indicated. ND, not determined.

Although the ∼50% efficiency observed in these experiments is more than sufficient to demonstrate the practical utility of the system, future systematic optimization of the protocol is likely to result in even greater overall efficiencies. In fact, during the first cycle of integration/excision, when the cargo of the pDASH-DI-ω1-MD7 was transferred to the pDASH-AIK, a second antibiotic was added to the media for the selection of the donor vector during rhamnose induction. This adjustment resulted in an increase in efficiency from ∼50% when using the standard procedure to ∼75% when using the double antibiotic selection (with 55%, 75%, and 90% efficiencies observed in three independent experiments). These encouraging results suggest that additional simple protocol optimizations, such as induction times, culture conditions, etc., could further improve the already high efficiency of the system.

### In planta functionality of DASH-generated constructs

To determine the functionality of the constructs generated using the DASH platform, we examined the activity of the genes present in the 97 kb pDASH-AIK-Big construct described above (Figure 5). Of the 35 transcriptional units in this construct (Figure 6A), *LacZ* and *sfGFP* should be functional in *E. coli*, while the rest should be active in plants. The functionality of *LacZ* and *sfGFP* was assessed by growing CZ105a cells carrying the pDASH-AIK-Big construct on LB plates supplemented with X-gal and by imaging the fluorescence of bacterial colonies under blue light, respectively. As can be observed in Figure 6B, both genes were active in *E. coli*. The rest of the 33 genes present in the pDASH-AIK-Big correspond to four identical copies of an eight transcriptional unit module and the *FAST* marker gene (Shimada *et al*., 2010). In this module, two different dCas9-EDLL activation domain fusion proteins (one from *Streptococcus pyogenic (Spy)* and another from *S. thermophilus* (*Sth*)), the corresponding *gRNA-MS2* fusions, and the MCP-VPR fusion protein are used to drive the expression of two reporter genes, *3xYpet* and *mCherry* (Figure 4). The expression of the *3xYPet* requires the activities of the transcriptional units TU1, 2, and 3, while the expression of the *mCherry* requires the activity of TU1, 5, and 7. In addition, the TU6 should confer kanamycin resistance in the corresponding transgenic plants (Figure 4), while the *FAST* transcriptional unit (Figure 5) should result in red fluorescent seed coats. Thus, to examine the functionality of these genes in plants, we first evaluated the activity of the *3xYpet* and *mCherry* in *N. benthamiana* transient expression assays. For that, the pDASH-AIK-Big was purified from CZ105a *E. coli* and electroporated into *Agrobacterium* GV3101. After selecting transformants in LB kanamycin plates and confirming the presence of the pDASH-AIK-Big construct by PCR using five pairs of diagnostic primers (Table S5), positive clones were grown in liquid media and used for agroinfiltration of *N. benthamiana* leaves (Zhang *et al*., 2020). Three days after infiltration, the activity of the two fluorescent proteins was examined using fluorescence microscopy. The expression of both 3xYPet and mCherry (Figure 6C) confirms the functionality of these two reporter genes and, therefore, that of the five other transcriptional units required for their activation. In addition, these experiments show the feasibility of using the pDASH-AIK acceptor vector in transient expression in *N. benthamiana*, even when the DNA cargos are very large, like in the current example.

**Figure 6.**
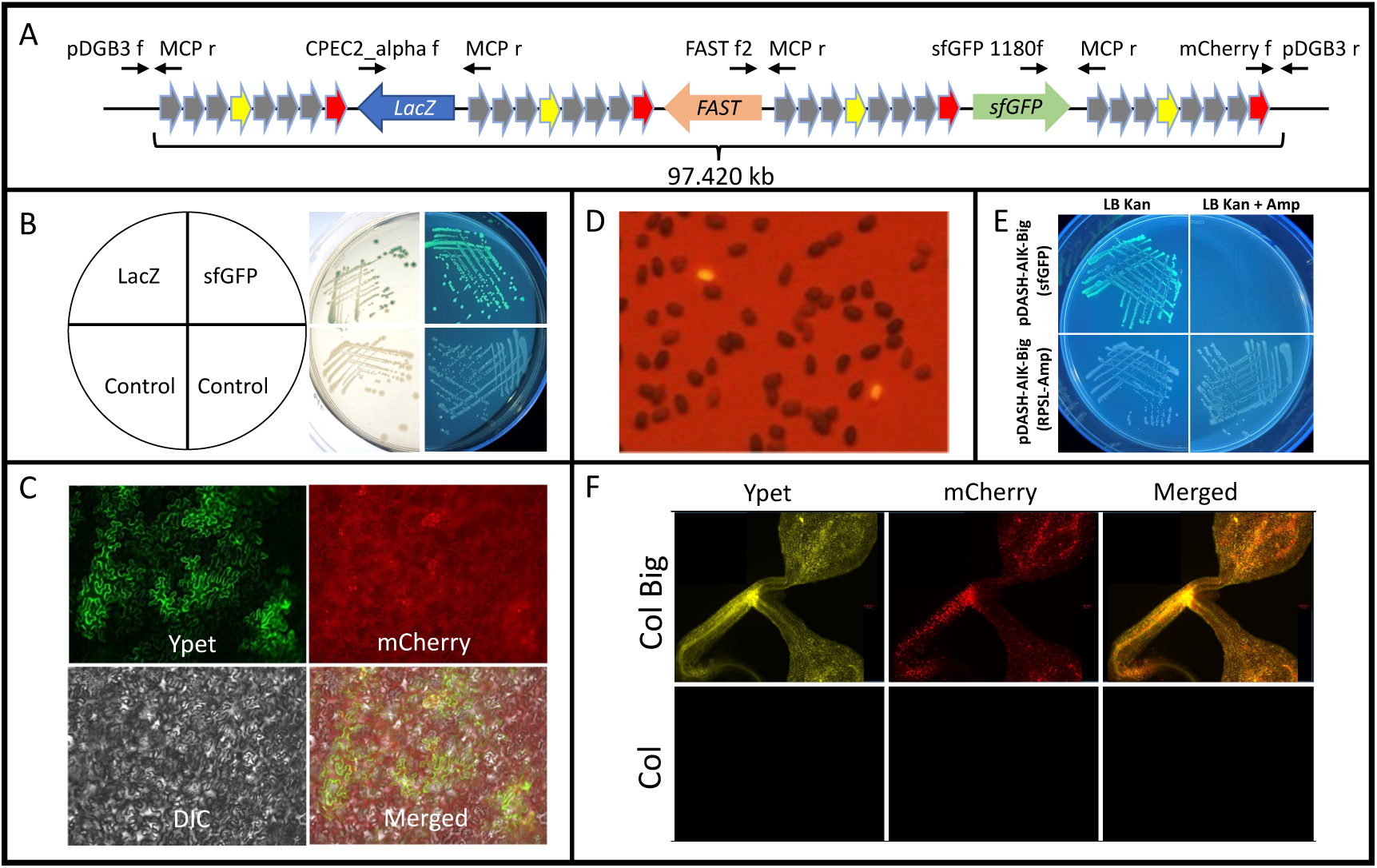
Functional characterization of the 116 kb construct (>97 kb T-DNA) generated with the DASH system. (A) Schematic representation of the final >97 kb cassette containing four repeats of Module 7 interspaced by the *lacZ*, *FAST* and *sfGFP* transcriptional units (pDASH-AIK-Big). Primers used to test the presence of the different components are shown. (B) Activity of the LacZ and sfGFP markers in bacteria harboring the pDASH-AIK-Big construct (upper) compared with the control (the acceptor vector only, pDASH-AIK) (lower). (C) Activity of the YPet and mCherry fluorescent proteins in *N. benthamiana* leaves transfected with *Agrobacterium* containing the pDASH-AIK-Big construct (enlarged cells shown in an inset). Expression of these two fluorescent proteins requires the activity of seven out of the eight transcriptional units comprising module 7. (D) Activity of the FAST reporter in seeds of Arabidopsis T0 plants. (E) Activity of the sfGFP fluorescence and Ampicillin-resistance markers in bacteria harboring the pDASH-AIK-Big construct before (top) and after (bottom) the recombineering-based post-assembly replacement of *sfGFP* with the *Ampicillin-*resistance gene. (F) Expression of the Ypet and mCherry in T1 Arabidopsis seedlings carrying the pDASH-AIK-Big T-DNA construct.

In addition to testing the functionality of the pDASH-AIK-Big in transient assays, we also generated Arabidopsis stable transgenic lines using the standard floral dip method (Clough and Bent, 1998). We first estimated the transformation efficiency by determining the percentage of red fluorescent seeds among the population of seeds from T0 plants (Figure 6D). The average efficiency of transformation in two independent experiments was much lower than when using standard binary vectors and just around 0.1%. These results agree with our previous observations using the original pYLTAC17 vector (Liu *et al*., 2002). We also screened the rest of the nonfluorescent seeds of the T0 plants for kanamycin resistance. We did not find any additional transformants, indicating that the FAST selection was effective, and the transformation efficiency estimated using this marker was accurate. We were also able to detect both Ypet and mCherry fluorescence in the transgenic plants obtained (Figure 6D and F). As with the transient expression in *N. benthamiana*, the detection of fluorescence indicates the functionality of the ∼97 kb T-DNA in stable transformants.

### Post-assembly modification capability of the DASH system

One of the key characteristics of the synthetic biology approach is its distinctive “design, build, test, learn” cycle. This engineering process often requires the redesign of specific DNA parts after evaluating their performance, often in the context of multigenic constructs. The development of a highly efficient assembly system greatly facilitates the process of generating these modifications by simply replacing one of the parts in a simple single-gene or small multigene assembly reaction. Redesigning constructs becomes considerably more challenging when dealing with large multigenic assemblies, as replacing a single component often requires restarting the entire assembly process, which typically involves multiple assembly cycles. Recombineering presents an attractive solution to this problem, as it allows for the introduction of any modifications (insertions, deletions, or replacements) in sequences of practically any size with high efficiency and precision. Thus, to facilitate the introduction of post-assembly modifications in the potentially large DASH constructs, the system has built-in recombineering capabilities. To demonstrate the functionality of this feature in the DASH system, we replaced the coding sequence of the *sfGFP* in the 97 kb cargo of pDASH-AIK-Big with the *RPSL-Amp* cassette sequences (Brumos *et al*., 2020). The 40 bp sequences flanking the *sfGFP* were incorporated at the ends of the *RPSL-Amp* recombineering cassette (Brumos *et al*., 2020) by PCR using the primers RPSL-Amp rep sfGFP f and RPSL-Amp rep sfGFP r. This PCR product was electroporated into recombineering-competent (Alonso and Stepanova, 2015) CZ105a cells carrying the 116 kb pDASH-AIK-Big construct. Ampicillin and kanamycin-resistant cells were selected in LB plates, and the replacement of *sfGFP* with *RPSL-Amp* was confirmed by PCR as well as functional assays (Figure 6 E, Table S5)

## Discussion

Manipulating gene expression using transgenic approaches is at the core of gene functional studies and a key element in biotechnological applications. The generation of the DNA constructs needed for these types of transgenic experiments can be significantly streamlined and accelerated using efficient synthetic biology DNA assembly tools. Among them, Golden Gate-based platforms have gained significant popularity due to their efficiency, experimental simplicity, and ability to utilize standardized, reusable DNA parts (Marillonnet and Grützner, 2020). One important limitation of these assembly technologies is that their efficiency greatly decreases as the size of the DNA parts being assembled increases, with a practical upper limit of about 25 kb, making the generation of relatively complex multigenic DNA constructs challenging (Lin and O’Callaghan, 2018). This could be a significant drawback as the demand for large DNA constructs to precisely control multigenic complex traits continues to increase in both synthetic biology and biotechnology applications. To specifically address this size limitation, highly efficient recombination-based gene stacking systems such as the GAANTRY and TGSII have been developed (Collier *et al*., 2018; Qin *et al*., 2022; Zhao *et al*., 2022; Zhu *et al*., 2017). While solving the size problem, the gene-stacking process in these systems imposes certain constraints. For instance, although the orientation and order of the genes in the final construct can be designed at will, once a gene is added, it cannot be removed, replaced, or modified without requiring a new assembly. Furthermore, none of these stacking systems are fully integrated with the popular Golden Gate-based platforms, resulting in a loss of the efficiency, standardization, and reusability benefits these platforms typically offer (Collier *et al*., 2018; Qin *et al*., 2022).

We developed the DASH system to overcome the shortcomings of existing DNA assembly and gene-stacking platforms while retaining their strengths. Thus, the DASH system combines the efficiency and experimental simplicity of the Golden Gate-based technologies with the large cargo capacity of recombination-based methods while offering the sequence modification flexibility characteristic of recombineering. To achieve this, we took advantage of a reengineered recombineering strain SW105, the GoldenBraid alpha and omega vectors, and the large-capacity TAC vector pYLTAC17. The SW105 *E. coli* recombineering strain has an arabinose-inducible *FLP* recombinase and a very tight temperature-inducible recombinase cassette comprising the *beta*, *gam*, and *exo* lambda phage genes. To enable this strain to perform unidirectional integration between two plasmids, we incorporated the PhiC31 integrase capability, which enables the unidirectional recombination between two heterotypic sites *in vivo* with high efficiency. Other platforms that also rely on similar integrases to catalyze the recombination between two plasmids *in vivo*, such as the GAANTRY, avoid possible undesirable effects associated with the expression of these recombinases by transiently expressing them from a helper vector (Collier *et al*., 2018). To minimize the number of elements in our system and, at the same time, ensure tight regulation in the expression of *PhiC3*1, we placed the coding sequence of this gene under the control of rhamnose-inducible operons in the *E. coli* genome of the SW105 recombineering strain. One potential issue with this strategy, compared to those that express the recombinase from a plasmid, is that, in our case, we use a single copy of *PhiC31* driven by an endogenous *E. coli* promoter. This raises the question of whether the expression levels will be sufficient to trigger efficient recombination between plasmids *in vivo*. With this potential problem in mind, we placed the *PhiC31* CDS under the control of two different endogenous rhamnose-inducible promoters, with the idea that at least one of them would provide a sufficient level of expression to trigger the integration but not so high as to result in toxic, detrimental effects. Of the two strains generated, CZ105a demonstrates normal growth and high efficiency of rhamnose-inducible PhiC31, making it a suitable choice for the DASH system.

We chose the GoldenBraid pDGB3 vectors (Vazquez-Vilar *et al*., 2017) as the foundation for the DASH system’s assembly system because they provide a straightforward and efficient method to combine sets of DNA parts and pairs of transcriptional units or modules reiteratively. Importantly, standardized DNA parts and assembled transcriptional units from various other Golden Gate-based platforms using *BsaI* as the cloning enzyme can be easily transferred to the DASH alpha donor vectors and, therefore, used in gene stacking reactions with the pDASH acceptor vector. Thus, for example, standardized DNA parts from GoldenBraid, Loop, Mobius, and MoClo, and assembled transcription units from GoldenBraid, Loop, Mobius, and Start-Stop can be directly transferred to a DASH alpha donor vectors. In addition, relatively simple modifications would be needed to create the DASH shuttle vectors or add compatibility linkers that allow for even greater integration of other type IIS restriction enzyme-based platforms, as we have done for MoClo (Figure 2B) and could be easily done in a similar fashion for other systems such as GreenGate (Piepers *et al*., 2023), JMC(Chamness *et al*., 2023), etc. (Figure 1). Finally, the developers of the GoldenBraid platform have continued to update the tools and expand the collection of plant parts [39] that our DASH system can also leverage.

Similarly, the pYLTAC17 vector was used as the backbone for the DASH acceptor vector due not only to its high-cargo capacity but also because it has been used extensively in recombineering experiments where plant genomic sequences harbored in a TAC are modified in *E. coli* and then transferred into plants via *Agrobacterium*-mediated transformation (Bitrián *et al*., 2011; Brumos *et al*., 2020; Zhou *et al*., 2011). Although the efficiency of plant transformation may not match the high levels achieved with standard binary vectors for small DNA constructs or the pRi *Agrobacterium* plasmid for larger constructs, it likely represents the best compromise when working with large DNA constructs that may require post-assembly modifications. This ability to propagate this binary vector in *E. coli* represents a significant advantage over the native pRi plasmid, which can only be propagated in *Agrobacterium*. Furthermore, we show that the pYLTAC17-derived pDASH-AIK vector can be used in *N. benthamiana* transient expression assays, broadening the downstream applications of the DASH system. Finally, although we have observed in the past a significant number of truncations of large T-DNAs inserted in the plant genome (Zhou *et al*., 2011), the simplicity by which two different selectable markers (kanamycin and FAST, for example) could be added at the beginning and end of the T-DNA should aid in the identification of plants carrying the complete insert.

As mentioned above, the basic DASH system consists of a single *E. coli* strain, four donor vectors, and one acceptor vector, reflecting the platform’s simplicity. We have developed seven additional vectors to facilitate specific experimental pipelines. Thus, for example, a second acceptor vector, pDASH-AIIK (Figure 2B), with an *attB^CC^* instead of the *attP^TT^* site, is also available, allowing the direct stacking of DII-type DASH donor vector and eliminating the need of an extra assembly reaction to move the cargo from a DII to a DI vector. Similarly, to assist in the removal of the donor vectors’ backbones after the integration and excision recombination reactions, a set of four donor vectors expressing the *SacB* counter-selectable marker gene has been generated (Figure 2B). SacB is toxic to cells grown in the presence of sucrose (Gay *et al*., 1985) and can be leveraged to get rid of the donor plasmid post-integration and backbone excision step, thus helping to bypass time-consuming plasmid re-transformation or re-streaking steps and colony-PCR screens. This eliminates the need to ‘curate’ the final construct through plasmid segregation by re-transforming, and instead involves simply growing the cells with the final construct in the presence of sucrose. Finally, to illustrate how simple modifications of the DASH system could be used to expand the compatibility of the DASH platform, the pDASH-DII-α1 (MoClo) and pDASH-DI-α2 (MoClo) were developed to allow for the transfer of transcriptional units from the MoClo level M plasmids to the DASH system.

One significant limitation of all current sequential DNA assembly and stacking systems is that any alteration, even a single nucleotide change, generally requires going back in the assembly chain and restarting from the point where the part necessitating modification was included. This obviously becomes especially critical in synthetic biology and modern plant biotechnological applications that often require relatively large multigenic constructs and several iterations of the “design, build, test, and learn” cycle. Recombineering represents an excellent post-assembly modification approach as it can deal with genome-size DNA molecules and generate variable-size insertions or deletions, from single nucleotide to several thousand base pairs (Alonso and Stepanova, 2015; Brumos *et al*., 2020). Obviously, any non-toxic DNA molecule that can be propagated in *E. coli,* and any recombineering-compatible bacterial strains, including *Agrobacterium* (Bian *et al*., 2022), can be modified by recombineering. Replacing *sfGFP* with an *RPSL-Amp* cassette in the ∼100 kb construct demonstrates that integrating recombineering into the DASH platform simplifies post-assembly modifications, even for very large DNA constructs.

Finally, the DASH system was developed with the idea of providing a user-friendly platform that, although capable of handling very large DNA molecules, does not require sophisticated laboratory techniques or users highly skilled in molecular biology or microbiology. The simplicity of the system is evident in the six elements that comprise it: the CZ105a *E. coli* strain, four donor vectors, and one acceptor vector. The simplicity of the experimental pipeline is demonstrated by the initial assembly of DNA components, which employs the classical Golden Gate-based single-tube restriction/ligation cloning reaction. Additionally, the stacking process requires only the preparation of competent cells harboring the acceptor vector, electroporation of the donor vector, induction of the desired recombinase upon the addition of the appropriate sugar, and selection of the recombinants with the proper antibiotic-supplemented media (Figure S5). We anticipate that the DASH system’s simplicity and ease of use, along with its high efficiency, robustness, and compatibility with various Golden Gate-based systems, combined with the increasing availability of GoldenBraid and other DASH-compatible tools and DNA parts, will make this platform appealing to a diverse group of plant biologists, particularly those who need to generate large DNA constructs or introduce post-assembly modifications.

## Materials and Methods

### Generation of PhiC31 integrase attachment site (*att*) and flippase recombinase target (*FRT*)

All *att* sites, including *attP^TT^*, AGTAGTGCCCCAACTGGGGTAACCT**TT**GAGTTCTCTCAGTTGGGGGCGTA, *attB^TT^*, CCGCGGTGCGGGTGCCAGGGCGTGCCC**TT**GGGCTCCCCGGGCGCGTACTCC, and their orthogonal *attP^CC^* (AGTAGTGCCCCAACTGGGGTAACCT**CC**GAGTTCTCTCAGTTGGGGGCGTA) and *attB^CC^* (CCGCGGTGCGGGTGCCAGGGCGTGCCC**CC**GGGCTCCCCGGGCGCGTACTCC) sites (Grindley *et al*., 2006; Merrick *et al*., 2018; Thorpe *et al*., 2000), *FRT*, GAAGTTCCTATACTTTCTAGAGAATAGGAACTTC (Senecoff *et al*., 1985), and combined *att* –*FRT* sites, were generated by PCR using primers listed in Table S1 and iProof High-Fidelity DNA Polymerase (www.bio-rad.com).

Briefly, to generate the *attP^TT^-FRT* site flanked by 40 bp of homology to each side of the desired insertion site in the pYLTAC17 (Liu *et al*., 2002) vector, we first amplified these flanking sequences using the primers P27/P83 and P86/P28 (see Table S1) and the pYLTAC17 vector as a template, and the *attP^TT^-FRT* site sequence was obtained by PCR with the overlapping primers P84/P85 (Table S1). The resulting three fragments were assembled together by Gibson assembly (www.neb.com) and cloned into pCR2.1 (www.thermofisher.com) according to the user manuals, generating pCR2.1_*attP^TT^-FRT* (acceptor).

Similarly, to obtain the *attB^CC^-FRT* sequence, the *FRT* sequence was amplified from pCR2.1_*attP^TT^-FRT* (acceptor) described above using primers P88/P70, while *attB^CC^* was generated with overlapping primers P69/P87. The resulting two fragments were assembled together via overlapping PCR with primers P69/P70 and cloned into pCR2.1, yielding pCR2.1_*attB^CC^-FRT* (omega1).

For the other seven *att* or *att-FRT* sites used to generate the different DASH donor vectors, a similar strategy was employed, utilizing either overlapping primers or amplifying the corresponding *att-FRT* plasmids as indicated in Table S1.

All PCR products were purified using QIAquick Gel Extraction Kit (www.qiagen.com) and cloned into pCR2.1 (www.thermofisher.com) according to the user manuals. All *att* and/or *FRT* sites in pCR2.1 were confirmed by Sanger sequencing.

## Donor vector modification

### Replacement of the kanamycin for the ampicillin resistance gene

The *kanamycin* gene of the GoldenBraid alpha-level vectors, pDGB3α1 and pDGB3α2 (Sarrion-Perdigones *et al*., 2011, 2013), was replaced with a domesticated *ampicillin* gene [(to remove an internal *BsaI* site, according to the GoldenBraid cloning (www.goldenbraidpro.com)] by recombineering (Alonso and Stepanova, 2015). Briefly, the domesticated *ampicillin* gene sequence was amplified using primers Amp rep Kan alpha f and r and the PCR product was used to replace the *kanamycin* sequences used standard recombineering procedures (Alonso and Stepanova, 2015). The resulting ampicillin version of pDGB3α1 and pDGB3α2, along with the original GoldenBraid omega-level vectors, pDGB3ω1 and pDGB3ω2, were utilized for subsequent donor vector modification.

### Insertion of the att and att-FRT sites in the donor vectors

To insert *att* and *att-FRT* just outside of the recognition sequence of a Type IIS restriction enzyme, *BsaI* or *BsmBI*, in the four donor vectors, the vectors were first linearized with restriction enzymes listed in Table S2. The backbones of all vectors and the *LacZ* cassettes for pDGB3ω1 and pDGB3ω2 were gel-purified. For pDGB3α1 and pDGB3α2 (ampicillin versions), since the *LacZ* cassettes contain restriction recognition sites for the restriction enzymes used to linearize the vectors, the intact *LacZ* cassettes were re-amplified from pDGB3α1 and pDGB3α2 by PCR with primers listed in Table S3. In addition, all *att* and *att*-*FRT* sites for each vector were also amplified from their corresponding pCR2.1-based plasmids listed in Table S1 by PCR with primers shown in Tables S1 and S3. Four DNA fragments (backbone, *LacZ* cassette, *att*, and *att-FRT*) for each donor vector were assembled by Gibson assembly (www.neb.com), generating pDASH-DII-α1, pDASH-DI-α2, pDASH-DI-ω1 and pDASH-DII-ω2.

### SacB domestication and insertion into donor vectors

The *SacB* gene was domesticated to remove the one *BsaI* and two *BsmBI* recognition sites with primers (P47-P54) listed in Table S3 from JMA1300 (Brumos *et al*., 2020) and cloned into pUPD2, according to GoldenBraid cloning. The domesticated *SacB* was inserted just downstream of the *ampicillin* gene in pDASH-DII-α1 and pDASH-DI-α2 or the *spectinomycin* gene in pDASH-DI-ω1 and pDASH-DII-ω2, forming an operon with the upstream antibiotic gene, by recombineering (Alonso and Stepanova, 2015), which was performed in two steps.

First, the *ampicillin* gene in donor alpha vectors or the *spectinomycin* gene in donor omega vectors was replaced with a *kanamycin*-*SacB* PCR fragment amplified with the primers listed in Table S3 by recombineering. In the second step, the *kanamycin* gene in all vectors was replaced with *ampicillin* in alpha or *spectinomycin* in omega vectors. For that, the *ampicillin* gene was first amplified using pDASH-DII-α1 DNA as a template and the primers Amp rep Kan alpha f and Amp rep Kan r2. The resulting PCR product was used to replace the *kanamycin* gene in the alpha vectors used standard recombineering procedures (Alonso and Stepanova, 2015), generating pDASH-DIIS-α1 and pDASH-DIS-α2.

Similarly, the primers Spec rep Kan f and r were used to amplify by PCR the *spectinomycin* sequences using DNA from the pDASH-DIS-ω1 vector and used in a recombineering experiment to replace the *kanamycin* sequences in the omega vectors. Due to the toxicity of *SacB* in the pDASH omega vectors, extra *spectinomycin* promoter sequences were removed by Gibson assembly (www.neb.com) using the fragments amplified by PCR with primers listed in Table S3, generating pDASH-DIS-ω1 or pDASH-DIIS-ω2.

### Generation of MoClo-compatible alpha donor vectors

The *LacZ* cassettes were amplified by PCR from GoldenBraid pDGB3α1 and pDGB3α2 with primers listed in Table S3, which introduced the MoClo-compatible assembly syntax. The resulting PCR products were purified, digested with *BsmBI*, and ligated with DASH donor vectors pDASH-DIIS-α1 and pDASH-DIS-α2 pre-digested with the same enzyme.

### Acceptor vector modification

The fragment containing *attP^TT^-FRT* with homology arms of 40 bp was amplified by PCR from the plasmid pCR2.1_*attP^TT^-FRT* (acceptor) with primers P27 and P28 (Table S1). The purified PCR products were used to replace the *SacB* cassette and the *RPSL-Amp* operon in the T-DNA region of JMA2450 by recombineering (Brumos *et al*., 2020), generating pDASH-AIK. For the second acceptor vector, pDASH-AIIK, plasmid pDASH-AIK-*LacZ* was digested by *BsaI*, filled in by *Klenow* (www.neb.com), and re-ligated by T4 DNA ligase. Both acceptor vectors were confirmed by whole plasmid sequencing and/or Sanger sequencing.

### *E. coli* SW105 strain modification for *PhiC31* expression under a rhamnose-inducible promoter

Two strains expressing *PhiC31* driven by a rhamnose-inducible promoter were made by recombineering (Alonso and Stepanova, 2015). Briefly, first, a positive selection and counterselection cassette, the *RPSL-Amp* operon in JMA1274 (Brumos *et al*., 2020), was amplified by PCR with primers, rhaTF and rhaTR, rhaA-Amp f and rhaA-Amp r, listed in Table S3. The resulting PCR products were purified and inserted immediately downstream of the *rhaT* gene or *rhaA* in the *rhaBAD* operon in the SW105 genome (Warming, 2005) through positive selection for ampicillin resistance using the standard recombineering procedures (Alonso and Stepanova, 2015). Second, *PhiC31* (its 605 aa version) was amplified from pET11*Phic31poly(A)* (Addgene plasmid #18942, (Groth *et al*., 2004)) by PCR with primers, phiC31 rhaF and phiC31 rhaR3, rhaA-phiC31 f and rhaA-phiC31 r, listed in Table S3. The resulting PCR products were used to replace the *RPSL-Amp* operon that had previously been inserted into the SW105 genome through counterselection for streptomycin resistance and standard recombineering procedures (Alonso and Stepanova, 2015), forming the *rhaT-PhiC31* or *rhaBA-PhiC31-D* operon in the genome (Figure S1). *PhiC31* and the corresponding flanking sequences in the *E. coli* genome were confirmed by Sanger sequencing.

## Construction of the donor vector constructs

### DNA parts/transcription units

The DNA parts, including *Arabidopsis thaliana U6-26* promoter and terminator (*AtU6p* and *AtU6t*) (Xing *et al*., 2014), *Solanum lycopersicum DFR* (*DIHYDROFLAVONOL-4-REDUCTASE*) gene promoter (*SlDFRp_Spy(SlDFR)_*), Cauliflower Mosaic Virus *35S* promoter (*35S*), *mCherry v1*, *3xYpet*, *3xNLS-EDLL*, and *Streptococcus pyogenes (Spy)* sgRNA2.1 targeting *SlDFRp_Spy(SlDRF)_* at the position –150 (*sgRNA_Spy(SlDFR)_-MS2*) (Selma *et al*., 2019), were directly amplified by PCR using iProof High-Fidelity DNA Polymerase (www.bio-rad.com) and subcloned into GoldenBraid entry vector, pUPD2. Templates and primers are listed in Table S4.

To generate the *Streptococcus thermophilus St1 (Sth) dCas9*-based CRISPRa transcriptional unit, the original protospacer and the *Spy* PAM site (GACTGGTTGGTGAGAGAAGAagg, where small letters indicate the *Spy* PAM site) at the position –150 of *SlDFRp_Spy(SlDFR)_* in pUPD2 were replaced by the *ADH1* protospacer followed by the *Sth* PAM sequence (GAAGTGGAGGTTGCTCCACCgcagaaa, where small letters indicate the *Sth* PAM site) (Steinert *et al*., 2015) via site-directed mutagenesis, according to QuickChange Site-Directed Mutagenesis kit’s manual (www.agilent.com), producing the pUPD2_*SlDFRp_Sth(ADH1)_* DNA part. Similarly, the Arabidopsis codon-optimized *Sth Cas9* orthologue was first amplified from pDE-*St1_Cas9* (Steinert *et al*., 2015) and subcloned into pUPD2, from which, dead *Sth Cas9*, *dCas9_Sth_* [*St1 dCas9* (D9A, H599A)] was generated by two rounds of inverse PCR with primers, StdCas9 D9A f and StdCas9 D9A r, StdCas9 H599A f and StdCas9 H599A r (Table S4). *Sth* engineered *sgRNA (v1)*-MS2 (Agudelo *et al*., 2020), *sgRNA_Sth(ADH1)_*-MS2, was created by overlapping PCR using primers, St gRNA scaffold f, Sth tracrRNA-short r, St gRNA ADH1f, Sth tracrRNA-short-MS2 f, and tracrRNA-MS2 75 r (Table S4).

To generate the pDASH-DII-α1_*FAST* transcriptional unit, the *FAST* cassette was domesticated from pDGE347_*FAST_Bar_pRPS5a:Cas9_ccdB* (Grützner *et al*., 2021) into the donor vector pDASH-DII-α1 according to GoldenBraid assembly to remove an internal *BsmBI* site. To generate the pDASH-DIIS-α1_*sfGFP*, the pDGB3alpha1_*AtU6p2*_*sfGFP* transcriptional unit (lab collection) was digested with *BsmBI*, releasing two fragments (backbone and AtU6p2_*sfGFP* cassette). The small fragment containing *sfGFP* cassette was purified and ligated with the DASH donor vector pDASH-DIIS-α1 pre-digested with the same enzyme. All sequences were confirmed by Sanger sequencing.

### Golden Gate Assembly

For the assembly of DNA parts into transcription units (TU) and individual TUs into modules, GoldenBraid assembly was performed according to NEB’s Golden Gate (24 fragment) Assembly Protocol (www.neb.com).

### PhiC31-mediated integration and FLP-mediated excision procedure

A single colony transformed with the acceptor (pDASH-AIK) and donor vectors was grown in 2 ml of liquid LB media supplemented with kanamycin and spectinomycin or ampicillin (depending on which donor vector was used) in a 15-ml sterile plastic culture tube at 30°C overnight. 20 µl of overnight culture were used to inoculate 1 ml of liquid LB media supplemented with antibiotics for the acceptor and donor vector selection. When OD_600_ reached ∼0.6, 10 µl of 10% sterile L-rhamnose were added to the cell culture to induce *PhiC31* and incubated for 3 hours at 30°C. *E. coli* cells were harvested by brief centrifugation and re-grown in 1 ml of liquid media supplemented with kanamycin and 10 µl of 10% sterile L-arabinose to induce *FLP* and incubated for 3 hours at 30°C. The culture was then re-streaked for single colonies on an LB plate supplemented with kanamycin. The plate was incubated at 30°C for 2 days for single colony selection and analysis.

### *Agrobacterium* and plant transformation

The pDASH-AIK-Big construct was transformed into *Agrobacterium tumefaciens* GV3101 by electroporation. After 2h incubation in plain LB at 28°C to allow the cells to recover and express the antibiotic-resistance genes, the electroporated cells were spread for single colonies on LB plates supplemented with kanamycin/gentamycin/rifamycin. Colonies were confirmed by colony PCR [by checking all junctions between two CRISPRa copies and *LacZ*, *FAST*, and *sfGFP*] with primers listed in Table S5 and used in transient expression assays in *N. benthamiana* leaves (by infiltrating the leaves with *Agrobacterium* suspension with OD_600_ of 0.1) (Beritza *et al*., 2024) or for Arabidopsis transformation via the floral dip method (Clough and Bent, 1998).

*N. benthamiana* (the laboratory accession, lab) seeds were sterilized using 50% (v/v) ethanol and directly sown onto soil (Sungro horticulture professional grow mix mixed 1:1 with Jolly gardener Pro-line C/B growing mix) for germination. At one-week-old stage, seedlings were transplanted to 24-cell nursery flats, one plant per cell, and grown at 22°C under a 16h-light/8h-dark cycle in a growth chamber with a light intensity of ∼120 μmol/m^2^/s.

Similarly, *Arabidopsis thaliana* (Col-0) seeds were sterilized using 50% (v/v) bleach solution and kept at 4°C for three days (for stratification), then directly sown onto soil described above with same growth conditions as for *N. benthamiana*.

### Microscopy to assess Ypet and mCherry fluorescence in plants

Samples were imaged using the Zeiss Axio Imager M2 upright microscope with a 5x objective (N.A. Immersion 0.16 and type Plan APO) and the corresponding multiple fluorescence channels, which were controlled by the Zeiss ZEN (blue edition) software. The Keyence BZ-X80 microscope was used with the 10x objective (N.A. Immersion 0.45 and type Plan APO). Three channels (Fluor GFP, Fluor TxRed, and BF/phase) were used to capture Ypet and mCherry fluorescence and bright field images, respectively. Images were exported as TIFF files.

## Author Contributions

Experiments were performed by CZ. Conceptualization, experimental design, and manuscript preparation were carried out by CZ, ANS, and JMA.

## Acknowledgements

This work was supported by the National Science Foundation grants 1940829, 1650139, and 1444561 to JMA and ANS, and 1750006 to ANS, and Research Capacity Fund (HATCH) project awards 7005468 and 7005482 from the U.S. Department of Agriculture’s National Institute of Food and Agriculture to JMA and ANS, respectively. We are grateful to Diego Orzaez’s group for sharing the GoldenBraid system, Dr. Holger Putchta for providing the *St1 Cas9* DNA, and Katie Vollen and Josefina-Patricia Fernandez-Moreno in the Alonso-Stepanova lab for sharing some GoldenBraid DNA parts.

---

**Figure S1.**
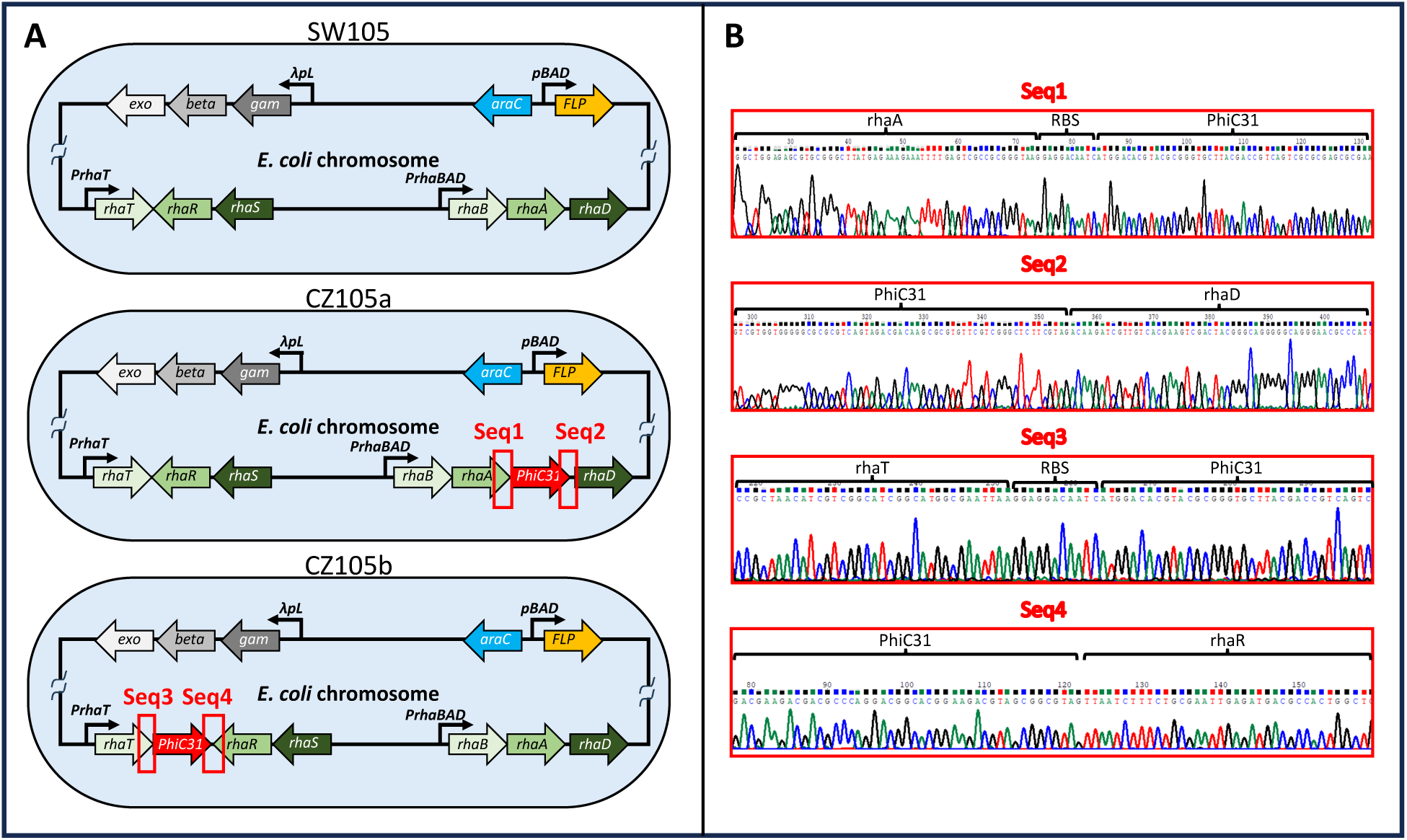
Genomic modifications in the DASH CZ105 strains. (A) Schematic representation of three strains of *E. coli*. The relevant genetic components of the original *E. coli* strain SW105, including a defective, temperature-sensitive λ prophage harboring three λ Red genes, *exo*, *beta*, and *gam,* for homologous recombination in the genome, are shown. In addition, the genome of the SW105 strain also carries an L-arabinose-inducible *FLP* gene cassette. To generate the DASH strain CZ105a, the phage-derived *PhiC31* integrase coding sequence and a ribosome binding site were inserted into the rhamnose-catabolism *rhaBAD* operon just downstream of the *rhaA* coding sequence. In the CZ105b strain, the same sequences were inserted just downstream of the transporter gene *rhaT*. (B) The Sanger sequencing chromatograms of the junctions between the *PhiC31* and the flanking sequencing in the *E. coli* genome are shown.

**Figure S2.**
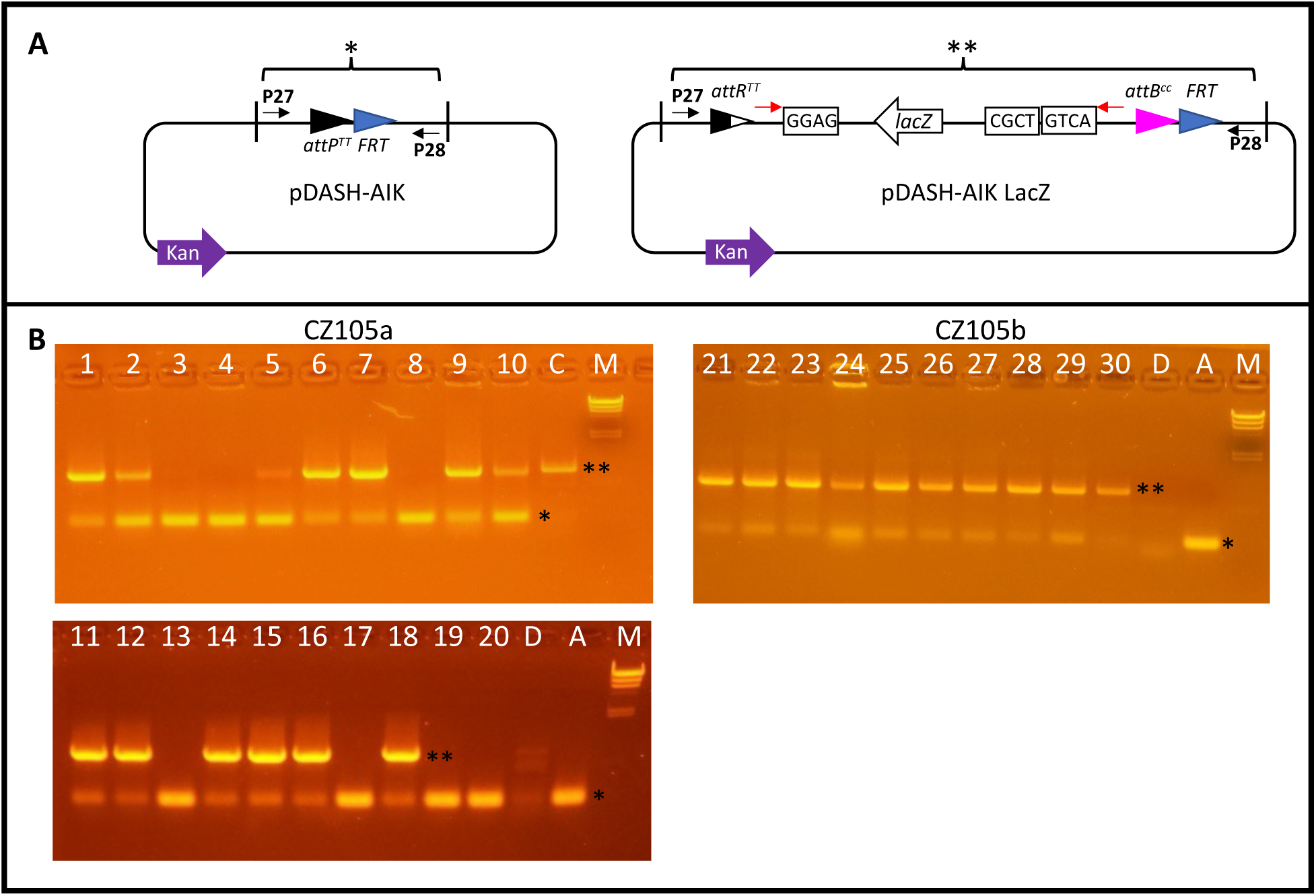
Functional analysis of the PhiC31 integrase and FLP recombinase activities in CZ105a and CZ105b cells. (A) Schematic representation of the pDASH-AIK acceptor vector before (left panel) and after (right panel) PhiC31-mediated integration and subsequent FLP-mediated excision of pDASH-DI-ω1 vector sequences. (B) PCR amplification of 20 CZ105a (left panels) and 10 CZ105b (right panel) colonies using the primers P27 and P28 flanking the recombination sites in the pDASH-AIK vector. The larger band (**) corresponds to plasmids where both the integration and excision have taken place, while the smaller band (*) corresponds to plasmids where no insertion has occurred. Whole plasmid sequencing of colony 30 indicates that weaker bands correspond to colonies where only a small fraction of the cells have undergone the integration and excision process. On the other hand, Sanger sequencing of the junction sites of the DNA from lane 7 indicates that strong bands correspond to colonies where integration and excision have both occurred. Letters M, D, and A indicate marker, donor, and acceptor DNA controls, respectively. Lane C in the CZ105a gels corresponds to colony 30 in gel CZ105b.

**Figure S3.**
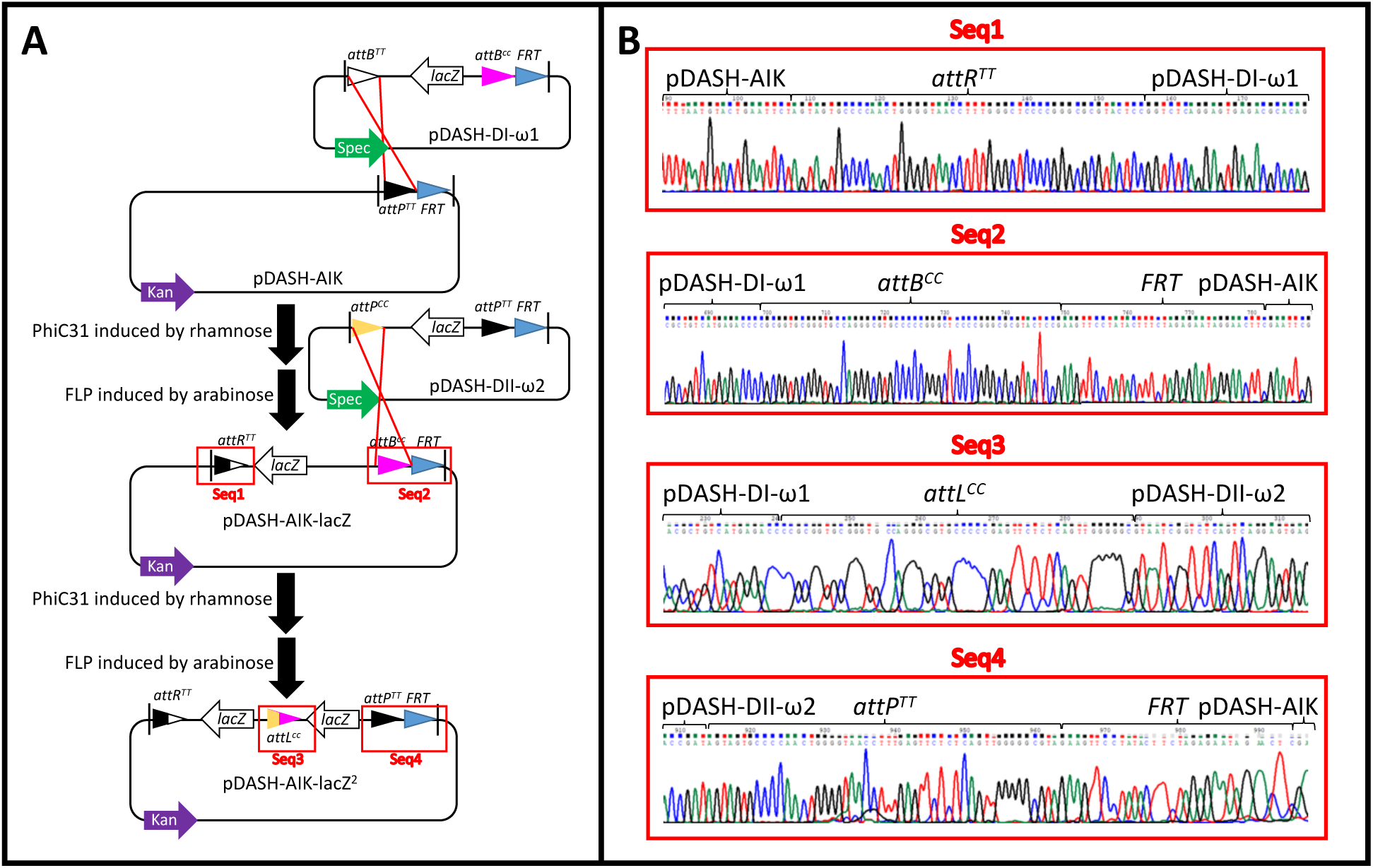
Fidelity of two consecutive rounds of PhiC31/FLP-mediated integration/excision reactions. (A) Schematic representation of the PhiC31/FLP-mediated integration and excision of pDASH-AIK and pDASH-DI-ω1 to generate the pDASH-AIK-lacZ construct, followed by a second round of integration and excision between the plasmids pDASH-AIK-lacZ and pDASH-DII-ω2 to produce the pDASH-AIK-lacZ^2^. (B) The chromatograms from the Sanger sequencing that correspond to the junction sites of the clones resulting from each integration/excision reaction, highlighted in red boxes in panel A, are displayed in panel B. Key parts of the sequences, such as recombination sites and marker genes, are annotated.

**Figure S4.**
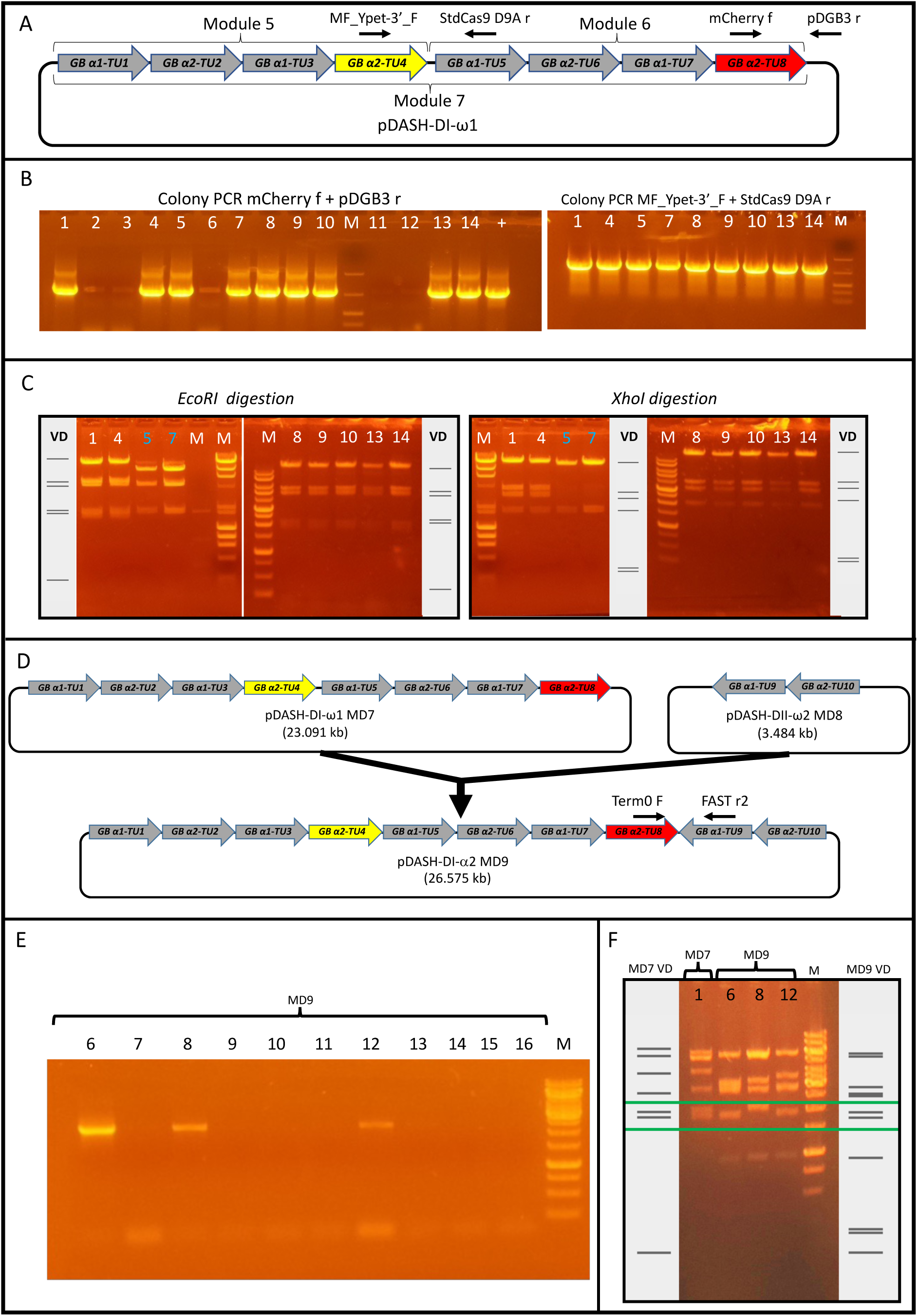
Example of the assembly efficiency estimation for large modules. (A) Schematic representation of module 7 obtained by assembling module 5 and module 6, as indicated by the brackets. The transcriptional units are depicted as large arrows, while the relative positions of the primers used for colony PCRs are marked with small black arrows. (B) The efficiency of assembling modules pDASH-DII-α1-MD5 and pDASH-DI-α2-MD6 to generate the 23 kb pDASH-DI-ω1-MD7 was determined by first examining the number of LacZ-positive (blue) colonies (not recombinant) and LacZ-negative (white) colonies (potential recombinants). Of the 51 colonies obtained, 14 were white, and the rest were blue. Colony PCR of the 14 white colonies using the diagnostic primers mCherry f and pDGB3 r indicates that 9 out of the 14 colonies contained sequences from Module 6 (left panel). PCRs with the primers MF_Ypet3’_F and StdCas9 D9Ar indicate that all these nine colonies contain sequences from Module 5 (right panel). (C) Diagnostic restriction digest of the plasmids corresponding to the nine PCR-positive colonies using two different enzymes, *EcoRI* or *XhoI*, indicates that seven of the nine colonies contain Module 7. Sequencing of the final 97 kb construct confirmed the fidelity of the assembly of Module 7. The incorrect clones based on the digestion pattern, 5 and 7, are marked with light blue numbers. M indicates a DNA marker, while “+” is a positive control. VD indicates virtual digestion. D) Schematic representation of module 9 attempted by assembling modules 7 and 8 together. The transcriptional units are depicted as large arrows, while the relative positions of the primers used for the colony PCRs are marked with small black arrows. (E) White colonies from module 9 assembly were examined by PCR using the primers Term0 F and FAST r2. Weak amplification was observed in 3 out of the 25 colonies examined (11 are shown). (F) Diagnostic restriction digest of the plasmids corresponding to three PCR-positive colonies with the enzyme *NsiI* indicates that none contain the intact module 9 sequence. Virtual digestion of module 7 (MD7 VD) and module 9 (MD9 VD) are shown. The green box highlights two diagnostic bands (2191 and 1994 bp in size) that should be identical in both MD7 and MD9 but are only seen in the MD7 control digestion (MD7 label) but not in any of the MD9 DNAs (MD9 label).

**Figure S5.**
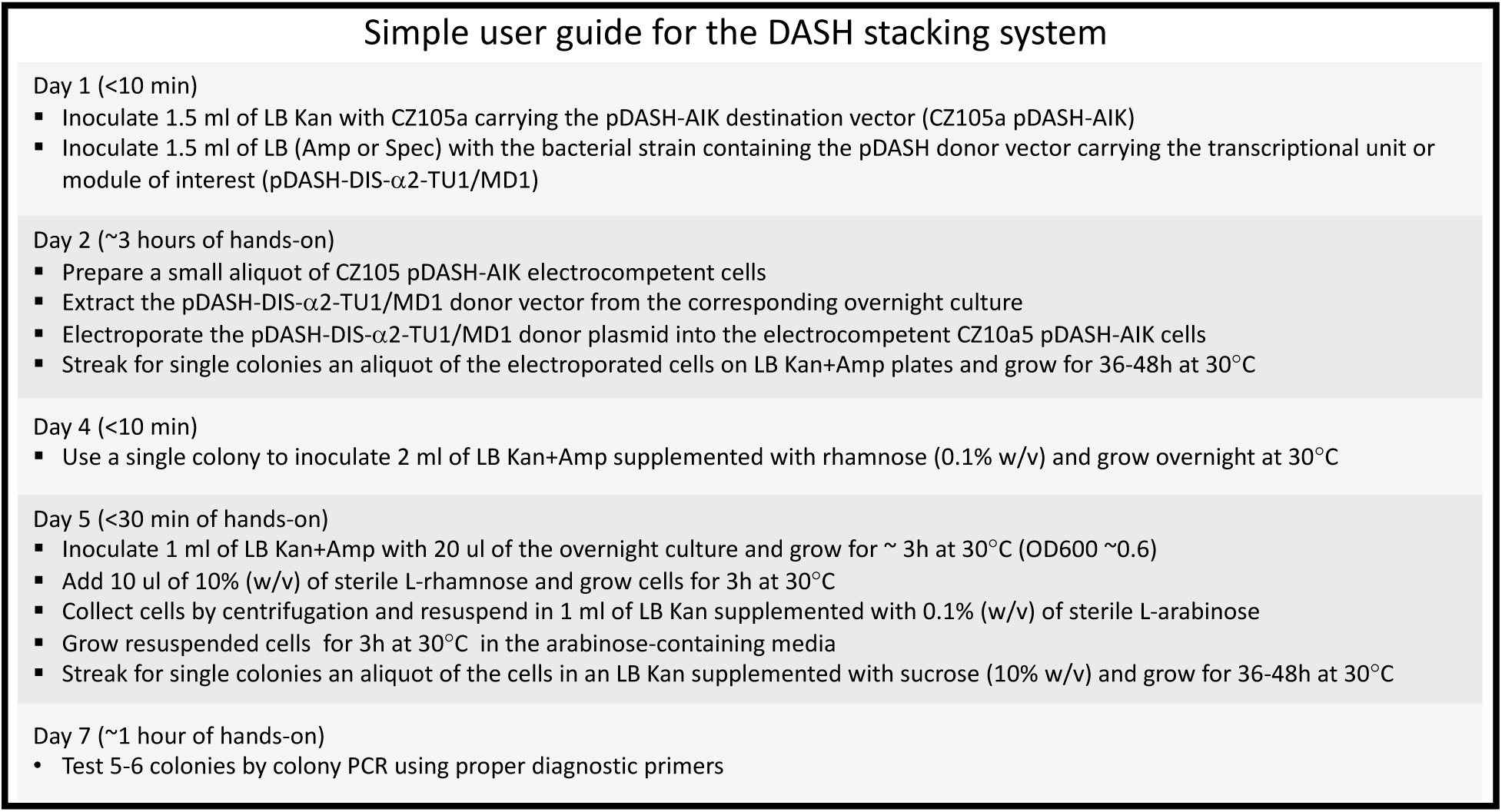
Simple user guide for the DASH gene stacking system. A brief overview of the steps involved in a typical gene stacking experiment using the DASH system is provided. The estimated hands-on time for each step and the total duration of the process are indicated. The guide assumes that the user already has the DASH destination vector in the CZ105a cells and that the transcriptional unit or module to be stacked is already present in a DASH donor vector.

**Table S1.**
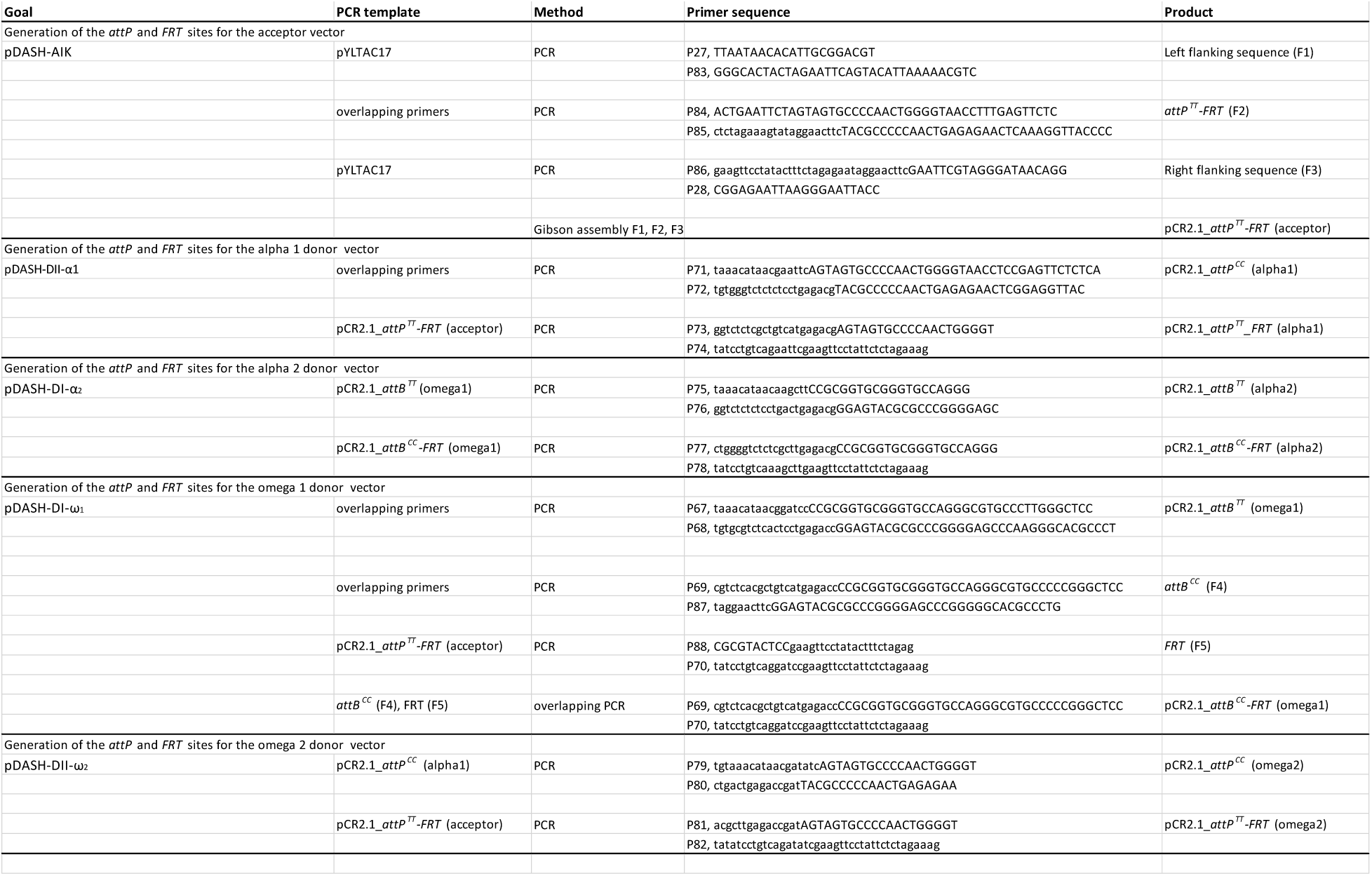
Incorporation of the PhiC31 integrase attachment site (*att*) and flippase recombinase target site (*FRT*) in DASH system vectors.

**Table S2.**
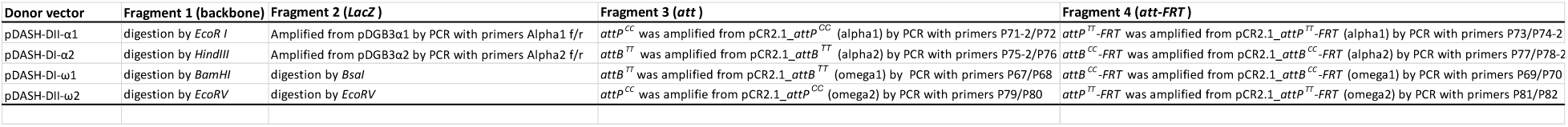
Incorporation of the *att* and *att-FRT* sites in the donor vectors.

**Table S3.**
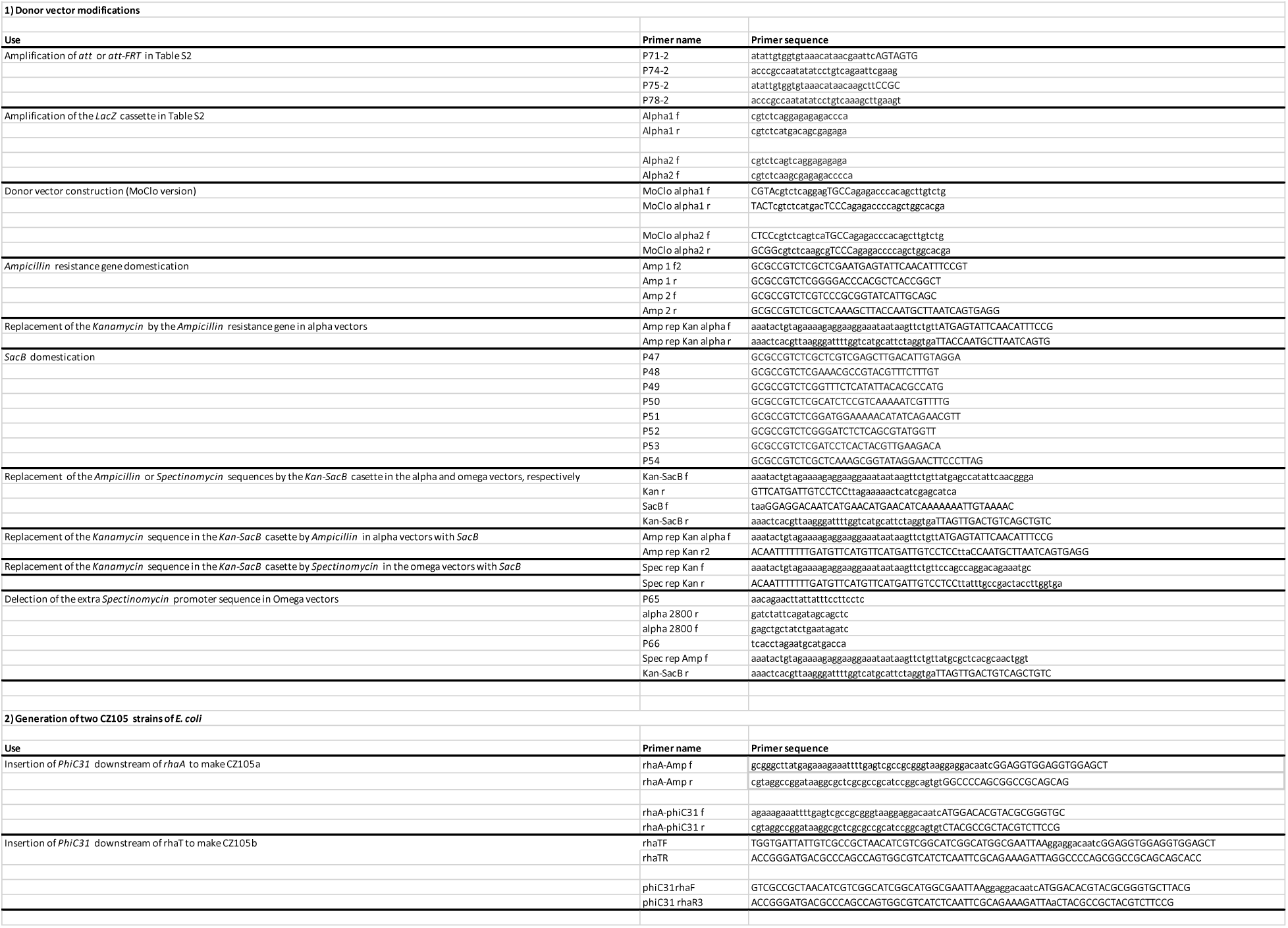
Primers used for the generation of the DASH donor vectors and the CZ105 *E. coli* strains.

**Table S4.**
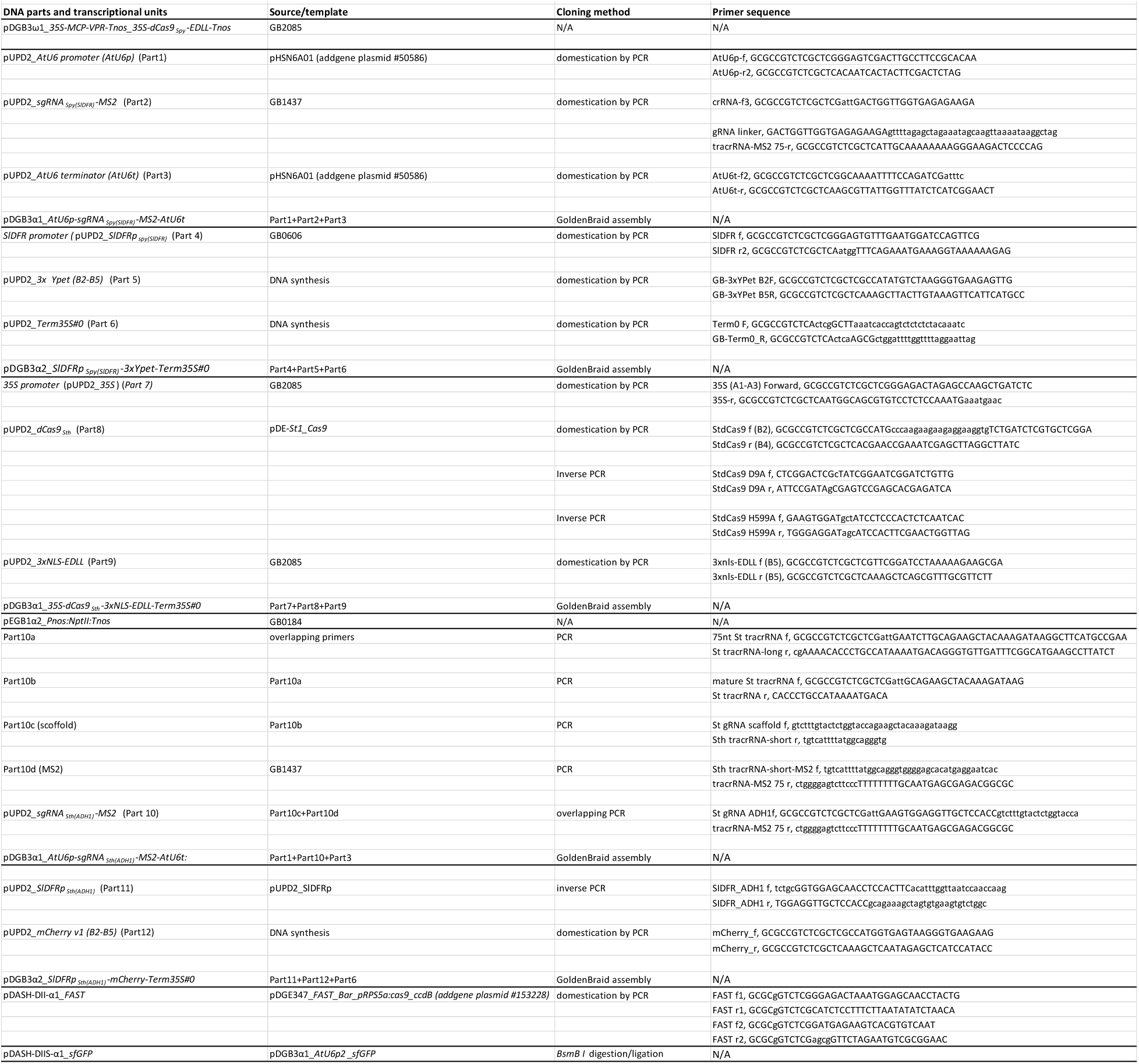
Generation of the donor vector constructs.

**Table S5.**
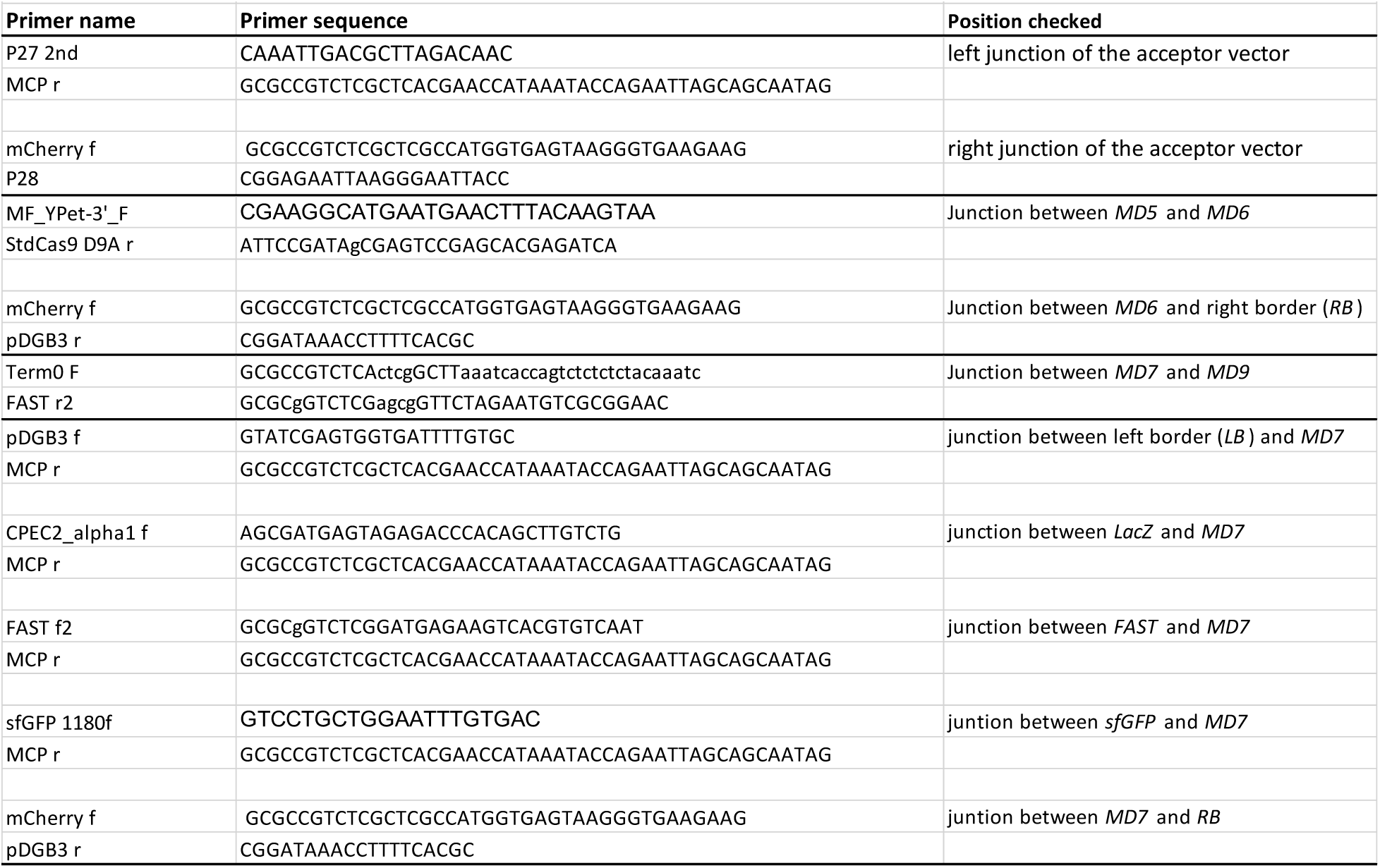
Primers used to check the integraty of the pDASH-AIK-Big construct.

## Notes

### Competing Interest Statement

The authors have declared no competing interest.

## References

1. Agudelo, D., Carter, S., Velimirovic, M., Duringer, A., Rivest, J.-F., Levesque, S., et al. (2020) *Versatile and robust genome editing with* Streptococcus thermophilus *CRISPR1-Cas9*. Genome Res., 30, 107– 117.

2. Ahrazem, O., Rubio-Moraga, A., Berman, J., Capell, T., Christou, P., Zhu, C., and Gómez-Gómez, L. (2016) The carotenoid cleavage dioxygenase CCD 2 catalysing the synthesis of crocetin in spring crocuses and saffron is a plastidial enzyme. New Phytologist, 209, 650–663.

3. Alonso, J.M. and Stepanova, A.N. (2015) A Recombineering-Based Gene Tagging System for Arabidopsis. In: Bacterial Artificial Chromosomes Methods in Molecular Biology (Narayanan, K., ed), pp. 233–243. New York, NY: Springer New York.

4. Andreou, A.I. and Nakayama, N. (2018) Mobius Assembly: A versatile Golden-Gate framework towards universal DNA assembly. PLoS ONE, 13, e0189892.

5. Andrews, B.J., Proteau, G.A., Beatty, L.G., and Sadowski, P.D. (1985) The FLP recombinase of the 2μ circle DNA of yeast: Interaction with its target sequences. Cell, 40, 795–803.

6. Beritza, K., Watts, E.C., and Van Der Hoorn, R.A.L. (2024) *Improving transient protein expression in agroinfiltrated* Nicotiana benthamiana. New Phytologist, 243, 846–850.

7. Bian, Z., Li, S., Yang, R., Yin, J., Zhang, Y., Tu, Q., et al. (2022) *Development of a New Recombineering System for* Agrobacterium *Species*. Appl Environ Microbiol, 88, e02499–21.

8. Bird, J.E., Marles-Wright, J., and Giachino, A. (2022) A User’s Guide to Golden Gate Cloning Methods and Standards. ACS Synth. Biol., 11, 3551–3563.

9. Bitrián, M., Roodbarkelari, F., Horváth, M., and Koncz, C. (2011) BAC-recombineering for studying plant gene regulation: developmental control and cellular localization of SnRK1 kinase subunits. The Plant Journal, 65, 829–842.

10. Blázquez, B., León, D.S., Torres-Bacete, J., Gómez-Luengo, Á., Kniewel, R., Martínez, I., et al. (2023) Golden Standard: a complete standard, portable, and interoperative MoClo tool for model and non-model proteobacteria. Nucleic Acids Research, 51, e98–e98.

11. Brumos, J., Zhao, C., Gong, Y., Soriano, D., Patel, A.P., Perez-Amador, M.A., et al. (2020) An Improved Recombineering Toolset for Plants. The Plant Cell, 32, 100–122.

12. Chamness, J.C., Kumar, J., Cruz, A.J., Rhuby, E., Holum, M.J., Cody, J.P., et al. (2023) An extensible vector toolkit and parts library for advanced engineering of plant genomes. The Plant Genome, 16, e20312.

13. Clough, S.J. and Bent, A.F. (1998) *Floral dip: a simplified method for* Agrobacterium *-mediated transformation of* Arabidopsis thaliana. The Plant Journal, 16, 735–743.

14. Collier, R., Thomson, J.G., and Thilmony, R. (2018) A versatile and robust Agrobacterium-based gene stacking system generates high-quality transgenic Arabidopsis plants. The Plant Journal, 95, 573–583.

15. De Paoli, H.C., Tuskan, G.A., and Yang, X. (2016) An innovative platform for quick and flexible joining of assorted DNA fragments. Sci Rep, 6, 19278.

16. Dusek, J., Plchova, H., Cerovska, N., Poborilova, Z., Navratil, O., Kratochvilova, K., et al. (2020) Extended Set of GoldenBraid Compatible Vectors for Fast Assembly of Multigenic Constructs and Their Use to Create Geminiviral Expression Vectors. Front. Plant Sci., 11, 522059.

17. Engler, C., Gruetzner, R., Kandzia, R., and Marillonnet, S. (2009) Golden Gate Shuffling: A One-Pot DNA Shuffling Method Based on Type IIs Restriction Enzymes. PLoS ONE, 4, e5553.

18. Engler, C., Kandzia, R., and Marillonnet, S. (2008) A One Pot, One Step, Precision Cloning Method with High Throughput Capability. PLoS ONE, 3, e3647.

19. Fricke, P.M., Gries, M.L., Mürköster, M., Höninger, M., Gätgens, J., Bott, M., and Polen, T. (2022) The l-rhamnose-dependent regulator RhaS and its target promoters from Escherichia coli expand the genetic toolkit for regulatable gene expression in the acetic acid bacterium Gluconobacter oxydans. Front. Microbiol., 13, 981767.

20. Gay, P., Le Coq, D., Steinmetz, M., Berkelman, T., and Kado, C.I. (1985) Positive selection procedure for entrapment of insertion sequence elements in gram-negative bacteria. J Bacteriol, 164, 918–921.

21. Gibson, D.G., Young, L., Chuang, R.-Y., Venter, J.C., Hutchison, C.A., and Smith, H.O. (2009) Enzymatic assembly of DNA molecules up to several hundred kilobases. Nat Methods, 6, 343–345.

22. Grindley, N.D.F., Whiteson, K.L., and Rice, P.A. (2006) Mechanisms of Site-Specific Recombination. Annu. Rev. Biochem., 75, 567–605.

23. Groth, A.C., Fish, M., Nusse, R., and Calos, M.P. (2004) Construction of Transgenic Drosophila by Using the Site-Specific Integrase From Phage φC31. Genetics, 166, 1775–1782.

24. Grützner, R., Martin, P., Horn, C., Mortensen, S., Cram, E.J., Lee-Parsons, C.W.T., et al. (2021) High-efficiency genome editing in plants mediated by a Cas9 gene containing multiple introns. Plant Communications, 2, 100135.

25. Hanko, E.K.R., Paiva, A.C., Jonczyk, M., Abbott, M., Minton, N.P., and Malys, N. (2020) A genome-wide approach for identification and characterisation of metabolite-inducible systems. Nat Commun, 11, 1213.

26. Hartley, J.L. (2000) DNA Cloning Using In Vitro Site-Specific Recombination. Genome Research, 10, 1788–1795.

27. Hirose, Y., Suda, K., Liu, Y., Sato, S., Nakamura, Yukino, Yokoyama, K., et al. (2015) *The Arabidopsis TAC Position Viewer: a high-resolution map of transformation-competent artificial chromosome (TAC) clones aligned with the* Arabidopsis thaliana *Columbia-0 genome*. The Plant Journal, 83, 1114–1122.

28. Kim, A.I., Ghosh, P., Aaron, M.A., Bibb, L.A., Jain, S., and Hatfull, G.F. (2003) *Mycobacteriophage Bxb1 integrates into the* Mycobacterium smegmatis groEL1 *gene*. Molecular Microbiology, 50, 463– 473.

29. Kirchmaier, S., Lust, K., and Wittbrodt, J. (2013) Golden GATEway Cloning – A Combinatorial Approach to Generate Fusion and Recombination Constructs. PLoS ONE, 8, e76117.

30. Lampropoulos, A., Sutikovic, Z., Wenzl, C., Maegele, I., Lohmann, J.U., and Forner, J. (2013) GreenGate – A Novel, Versatile, and Efficient Cloning System for Plant Transgenesis. PLoS ONE, 8, e83043.

31. Lee, E.-C., Yu, D., Martinez De Velasco, J., Tessarollo, L., Swing, D.A., Court, D.L., et al. (2001) A Highly Efficient Escherichia coli-Based Chromosome Engineering System Adapted for Recombinogenic Targeting and Subcloning of BAC DNA. Genomics, 73, 56–65.

32. Lin, D. and O’Callaghan, C.A. (2018) MetClo: methylase-assisted hierarchical DNA assembly using a single type IIS restriction enzyme. Nucleic Acids Research.

33. Liu, Y.-G., Liu, H., Chen, L., Qiu, W., Zhang, Q., Wu, H., et al. (2002) Development of new transformation-competent artificial chromosome vectors and rice genomic libraries for efficient gene cloning. Gene, 282, 247–255.

34. Marillonnet, S. and Grützner, R. (2020) Synthetic DNA Assembly Using Golden Gate Cloning and the Hierarchical Modular Cloning Pipeline. CP Molecular Biology, 130, e115.

35. Merrick, C.A., Zhao, J., and Rosser, S.J. (2018) Serine Integrases: Advancing Synthetic Biology. ACS Synth. Biol., 7, 299–310.

36. Moralejo, P., Egan, S.M., Hidalgo, E., and Aguilar, J. (1993) Sequencing and characterization of a gene cluster encoding the enzymes for L-rhamnose metabolism in Escherichia coli. J Bacteriol, 175, 5585–5594.

37. Park, J., Throop, A.L., and LaBaer, J. (2015) Site-Specific Recombinational Cloning Using Gateway and In-Fusion Cloning Schemes. *CP Molecular Biology*, 110.

38. Piepers, M., Erbstein, K., Reyes-Hernandez, J., Song, C., Tessi, T., Petrasiunaite, V., et al. (2023) GreenGate 2.0: Backwards compatible addons for assembly of complex transcriptional units and their stacking with GreenGate. PLoS ONE, 18, e0290097.

39. Pollak, B., Cerda, A., Delmans, M., Álamos, S., Moyano, T., West, A., et al. (2019) Loop assembly: a simple and open system for recursive fabrication of DNA circuits. New Phytologist, 222, 628–640.

40. Pollak, B., Matute, T., Nuñez, I., Cerda, A., Lopez, C., Vargas, V., et al. (2020) Universal loop assembly: open, efficient and cross-kingdom DNA fabrication. Synthetic Biology, 5, ysaa001.

41. Qin, G., Wu, S., Zhang, L., Li, Y., Liu, C., Yu, J., et al. (2022) An Efficient Modular Gateway Recombinase-Based Gene Stacking System for Generating Multi-Trait Transgenic Plants. Plants, 11, 488.

42. Sarrion-Perdigones, A., Falconi, E.E., Zandalinas, S.I., Juárez, P., Fernández-del-Carmen, A., Granell, A., and Orzaez, D. (2011) GoldenBraid: an iterative cloning system for standardized assembly of reusable genetic modules. PLoS One, 6, e21622.

43. Sarrion-Perdigones, A., Vazquez-Vilar, M., Palaci, J., Castelijns, B., Forment, J., Ziarsolo, P., et al. (2013) GoldenBraid 2.0: A Comprehensive DNA Assembly Framework for Plant Synthetic Biology. PLANT PHYSIOLOGY, 162, 1618–1631.

44. Sasaki, Y., Sone, T., Yoshida, S., Yahata, K., Hotta, J., Chesnut, J.D., et al. (2004) Evidence for high specificity and efficiency of multiple recombination signals in mixed DNA cloning by the Multisite Gateway system. Journal of Biotechnology, 107, 233–243.

45. Selma, S., Bernabé-Orts, J.M., Vazquez-Vilar, M., Diego-Martin, B., Ajenjo, M., Garcia-Carpintero, V., et al. (2019) Strong gene activation in plants with genome-wide specificity using a new orthogonal CRISPR /Cas9-based programmable transcriptional activator. Plant Biotechnology Journal, 17, 1703–1705.

46. Senecoff, J.F., Bruckner, R.C., and Cox, M.M. (1985) The FLP recombinase of the yeast 2-micron plasmid: characterization of its recombination site. Proc. Natl. Acad. Sci. U.S.A., 82, 7270–7274.

47. Shih, P.M., Vuu, K., Mansoori, N., Ayad, L., Louie, K.B., Bowen, B.P., et al. (2016) A robust gene-stacking method utilizing yeast assembly for plant synthetic biology. Nat Commun, 7, 13215.

48. Shimada, T.L., Shimada, T., and Hara-Nishimura, I. (2010) *A rapid and non-destructive screenable marker, FAST, for identifying transformed seeds of* Arabidopsis thaliana. The Plant Journal, 61, 519–528.

49. Steinert, J., Schiml, S., Fauser, F., and Puchta, H. (2015) *Highly efficient heritable plant genome engineering using Cas9 orthologues from* Streptococcus thermophilus *and* Staphylococcus aureus. The Plant Journal, 84, 1295–1305.

50. Sternberg, N., Sauer, B., Hoess, R., and Abremski, K. (1986) Bacteriophage P1 cre gene and its regulatory region. Journal of Molecular Biology, 187, 197–212.

51. Szybalski, W., Kim, S.C., Hasan, N., and Podhajska, A.J. (1991) Class-IIS restriction enzymes — a review. Gene, 100, 13–26.

52. Tao, Q. (1998) Cloning and stable maintenance of DNA fragments over 300 kb in Escherichia coli with conventional plasmid-based vectors. Nucleic Acids Research, 26, 4901–4909.

53. Taylor, G.M., Mordaka, P.M., and Heap, J.T. (2019) Start-Stop Assembly: a functionally scarless DNA assembly system optimized for metabolic engineering. Nucleic Acids Research, 47, e17–e17.

54. Thorpe, H.M. and Smith, M.C.M. (1998) In vitro *site-specific integration of bacteriophage DNA catalyzed by a recombinase of the resolvase/invertase family*. Proc. Natl. Acad. Sci. U.S.A., 95, 5505–5510.

55. Thorpe, H.M., Wilson, S.E., and Smith, M.C.M. (2000) *Control of directionality in the site-specific recombination system of the* Streptomyces *phage φC31*. Molecular Microbiology, 38, 232–241.

56. Vazquez-Vilar, M., Quijano-Rubio, A., Fernandez-del-Carmen, A., Sarrion-Perdigones, A., Ochoa-Fernandez, R., Ziarsolo, P., et al. (2017) *GB3.0: a platform for plant bio-design that connects functional DNA elements with associated biological data*. *Nucleic Acids Res*, gkw1326.

57. Warming, S. (2005) Simple and highly efficient BAC recombineering using galK selection. Nucleic Acids Research, 33, e36–e36.

58. Weber, E., Engler, C., Gruetzner, R., Werner, S., and Marillonnet, S. (2011) A Modular Cloning System for Standardized Assembly of Multigene Constructs. PLoS ONE, 6, e16765.

59. Xing, H.-L., Dong, L., Wang, Z.-P., Zhang, H.-Y., Han, C.-Y., Liu, B., et al. (2014) A CRISPR/Cas9 toolkit for multiplex genome editing in plants. BMC Plant Biol, 14, 327.

60. Yu, D., Ellis, H.M., Lee, E.-C., Jenkins, N.A., Copeland, N.G., and Court, D.L. (2000) *An efficient recombination system for chromosome engineering in* Escherichia coli. Proc. Natl. Acad. Sci. U.S.A., 97, 5978–5983.

61. Zhang, Y., Buchholz, F., Muyrers, J.P.P., and Stewart, A.F. (1998) A new logic for DNA engineering using recombination in Escherichia coli. Nat Genet, 20, 123–128.

62. Zhang, Y., Chen, M., Siemiatkowska, B., Toleco, M.R., Jing, Y., Strotmann, V., et al. (2020) A Highly Efficient Agrobacterium-Mediated Method for Transient Gene Expression and Functional Studies in Multiple Plant Species. Plant Communications, 1, 100028.

63. Zhao, Y., Han, J., Tan, J., Yang, Y., Li, S., Gou, Y., et al. (2022) *Efficient assembly of long DNA fragments and multiple genes with improved nickase-based cloning and Cre/* LOXP *recombination*. Plant Biotechnology Journal, 20, 1983–1995.

64. Zhou, R., Benavente, L.M., Stepanova, A.N., and Alonso, J.M. (2011) A recombineering-based gene tagging system for Arabidopsis. The Plant Journal, 66, 712–723.

65. Zhu, Q., Yu, S., Zeng, D., Liu, H., Wang, H., Yang, Z., et al. (2017) Development of “Purple Endosperm Rice” by Engineering Anthocyanin Biosynthesis in the Endosperm with a High-Efficiency Transgene Stacking System. Molecular Plant, 10, 918–929.

